# Multi-omics characterization of IDH-mutant astrocytoma-derived cell lines reveals NOTCH-regulated plastic quiescent astrocyte-like state

**DOI:** 10.64898/2025.12.30.696808

**Authors:** L. Garcia, C. Granotier-Beckers, D. Pineau, D. Sika, S. Hideg, C. Labadie, K. Aguilar Cázarez, J. Kundnani, L. Stuani, L. Gauthier, F. Boussin, M-A. Mouthon, S. Urbach, K. El Koulali, M. Séveno, C. Ripoll, K. Daddi, M. Verreault, A. Idbaih, S. O’Connor, C. Plaisier, L. Zhang, M. Zheng, V. Rigau, H. Duffau, J-P. Hugnot

**Affiliations:** Institut de Génomique Fonctionnelle (IGF), CNRS, INSERM, Université de Montpellier, Montpellier, France; Université Paris Cité, Inserm, CEA, Stabilité Génétique Cellules Souches et Radiations, LRP/iRCM/IBFJ, F-92265, Fontenay-aux-Roses, France; Université Paris-Saclay, Inserm, CEA, Stabilité Génétique Cellules Souches et Radiations, LRP/iRCM/IBFJ, F-92265, Fontenay-aux-Roses, France; Institut de Génétique Humaine (IGH), CNRS, Montpellier, France; Université de Montpellier, France; Institut de Recherche en Cancérologie (IRCM), INSERM, Montpellier, France; Equipe labélisée Ligue Contre le Cancer, Paris, France; BCM, Université Montpellier, CNRS, INSERM, Montpellier, France; Sorbonne Université, AP-HP, Institut du Cerveau (ICM), CNRS, INSERM, Hôpitaux Universitaires La Pitié Salpêtrière - Charles Foix, Service de Neuro-Oncologie-Institut de Neurologie, F-75013, Paris, France; School of Biological and Health Systems Engineering, Arizona State University, Tempe AZ, USA; Jinfeng Laboratory, Chongqing, China; CHU-Gui de Chauliac, Montpellier, France

## Abstract

Diffuse IDH-mutant astrocytomas are brain tumors typically diagnosed as low-grade but capable of progressing to higher grades. They exhibit three cellular states resembling astrocytes, oligodendrocytes, and neural progenitor (NPC) cells. Understanding their biology has been challenging due to the lack of relevant in vitro models.

Here we established and extensively characterized four astrocytoma cell lines (LGG275, LGG336, LGG85, LGG349) derived from IDH-mutant astrocytoma at different grades, cultured in defined media and analyzed by multi-omics. These lines display growth rates *in vitro* and *in vivo* consistent with tumor grade and recapitulate key molecular alterations observed in patient tumors, including *IDH1*, *ATRX*, and *TP53* mutations, activation of the alternative lengthening of telomeres (ALT) pathway and, in the most aggressive line, amplification of *MET* and *PDGFRA*.

Single-cell RNA sequencing showed that the 4 astrocytoma lines maintain the three major cellular states observed in patient tumors. A hallmark of higher-grade-derived lines (LGG85, LGG349) is the persistence of NPC-like populations without growth factors, reflecting tumor progression. The LGG275 line most accurately mirrors slow-growing astrocytomas. Using CD44 and GLAST, we isolated astrocyte-like (CD44+/GLAST+) cells from LGG275 that preferentially adopt a quiescent state yet retain remarkable plasticity, generating oligodendrocyte-like cells (CD44−/GLAST−). Transcriptomic and proteomic analyses revealed that astrocyte-like and oligodendrocyte-like cells populations resemble quiescent and activated neural stem (NSC) cells from the adult subventricular zone (SVZ).

Finally, we found that NOTCH signaling regulates the balance between astrocytic and oligodendrocytic states, while DLL3, expressed by oligodendrocyte-like cells, modulates both proliferation and phenotype. These cell lines represent valuable resources for dissecting lineage dynamics, heterogeneity, and progression mechanisms in IDH-mutant astrocytomas.

**Graphical Abstract:** 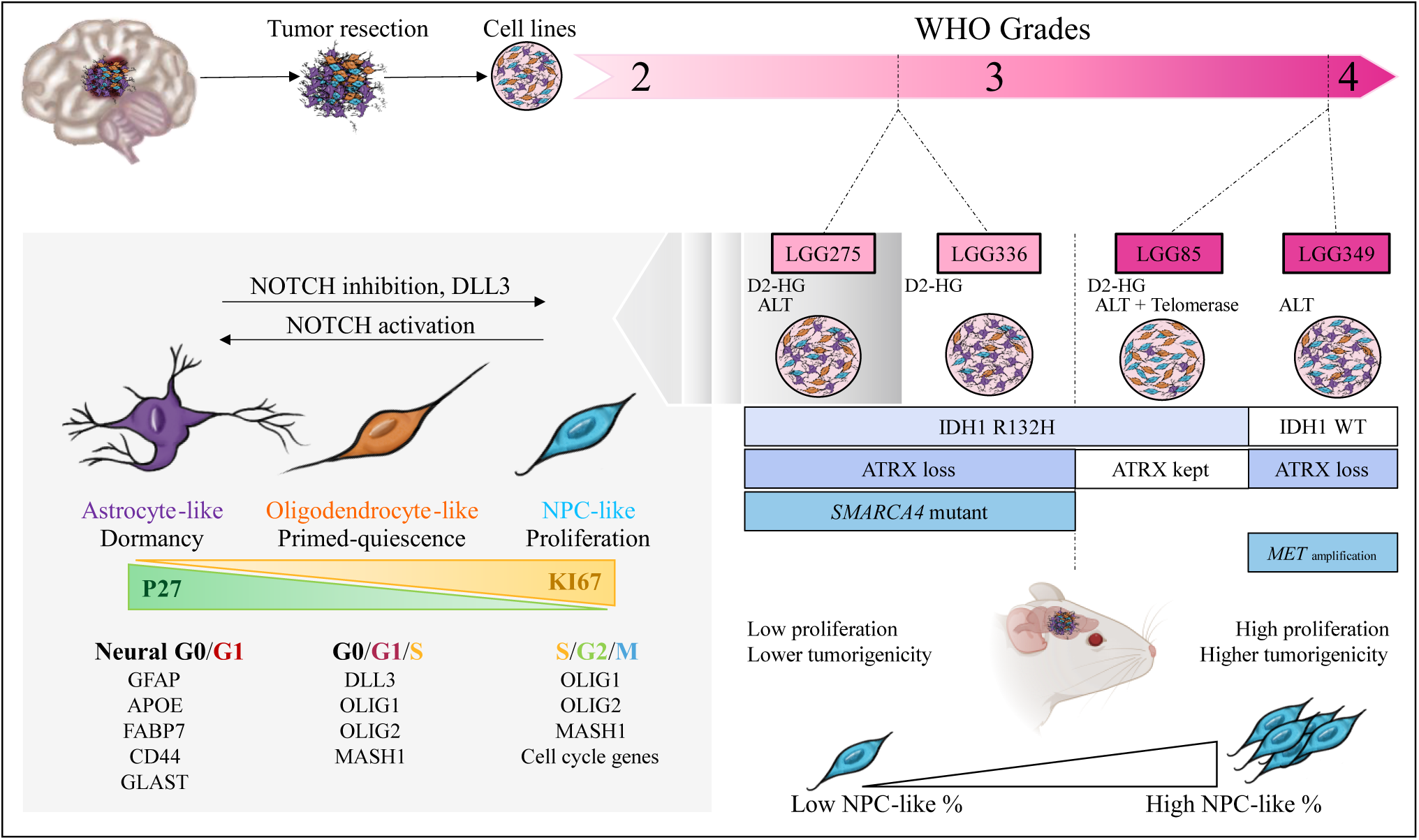

**Highlight:**

- We constituted a richly annotated biobank derived from 4 astrocytomas, showing similar characteristics to those found in patients, providing valuable tools to investigate cellular heterogeneity and plasticity, and link with tumor progression.
- scRNA-sequencing revealed three cell states (oligodendrocyte-like, astrocyte-like, and stem cell-like cells) akin to those found in tumors.
- Astrocyte-like cells are quiescent cells, plastic and similar to quiescent neural stem cell (qNSC) from the sub-ventricular zone (SVZ), while oligodendrocyte-like cells are similar to active NSC (aNSC).
- The Notch pathway plays a role in cell plasticity, enabling a shift towards an astrocyte-like state.

## Introduction

Diffuse low-grade gliomas (DLGG) are rare primary brain tumors that mainly affect young adults[1]. Molecularly, approximately 80% of DLGG harbor a missense mutation in *IDH1* or *IDH2*, leading to the production of the oncometabolite 2-hydroxyglutarate (2-HG)[2], which acts as an epigenetic disruptor, impairing cellular differentiation[3]. According to the World Health Organization (WHO) classification[4]: two major IDH1-mutant subtypes are recognized: astrocytomas, characterized by ATRX and TP53 mutations and exhibiting an alternative lengthening of telomeres (ALT) phenotype[5], and oligodendrogliomas, defined by 1p/19q co-deletion, together with CIC and FUBP1 alterations. These tumors, most often diagnosed as CNS WHO grade 2, are typically slow-growing (≈ 3-5 mm/year in diameter), with a low proliferative index (KI67 usually < 5%). However, they can evolve into more aggressive forms corresponding to grades 3 and 4. This malignant progression is driven by the acquisition of additional genetic alterations, such as *CDKN2A* deletion, *SMARCA4* mutations, and *PDGFRA* or *MET* amplifications, all of which are associated with a poorer prognosis[6].

Like glioblastomas, DLGG are intrinsically heterogeneous at the molecular and cellular levels. We and others have shown that these tumors comprise three major cellular states: astrocyte-like cells, oligodendrocyte-like cells and proliferative neural progenitor-like (NPC-like) cells[7–9]. These populations closely resemble their counterparts in the normal brain; astrocytes, oligodendrocyte precursor cells (OPC) and neural stem/progenitor cells, and express corresponding astrocytic identity, oligodendroglial lineage and markers of stemness/proliferation, respectively. Notably, only the NPC-like fraction is actively dividing, representing less than 10% of tumor cells, and this proportion increases with tumor grade[8,9]. In addition, as in many other tumors, gliomas display heterogeneity in cell-cycle dynamics, with subsets of cells remaining quiescent while others are actively engaged in various stages of the cell cycle[10]. Several studies have highlighted that quiescent tumor cells are more resistant to therapy and likely contribute to glioma recurrence following treatment[11].

The study of IDH-mutant astrocytomas has long been constrained by the scarcity of relevant *in vitro* models, limiting our ability to investigate their biological features and malignant progression mechanisms. To date, only few astrocytoma cell lines have been reported and well characterized[12,13], underscoring the need for new experimental systems that faithfully recapitulate these tumors. To address this gap, in this study, we established and thoroughly characterized four novel cell lines derived from IDH-mutant astrocytomas of different grades to provide cellular tools for mechanistic and translational research. We sought to determine whether astrocyte-like, oligodendrocyte-like, and NPC-like states are maintained within these cultures, to explore their relationships with quiescence and plasticity, and to identify some molecular determinants governing state transitions. Furthermore, we sought to find markers to purify and study astrocyte-like sub populations and study their properties. Together, these models offer valuable tools and molecular resources to investigate lineage dynamics, cellular heterogeneity, and the mechanisms driving tumor progression in IDH1-mutant astrocytomas.

## Results

### Isolation and characterization of 4 cell lines isolated from IDH1-mutant astrocytomas

Using explant cultures maintained in defined mediums supplemented with EGF and FGF_2_, we successfully established four adherent cell lines from patients diagnosed with IDH1-mutant gliomas (**Fig.1A**). Cell lines were considered established upon successful maintenance beyond 10 passages[7,14,15]. LGG275[7] and LGG336 were derived from histologically classified grade 2 astrocytomas, whereas LGG85[15] and LGG349 originated grade 4 astrocytomas (**Table.S1**). The LGG275 patient’s molecular analysis showed a hemizygous *CDKN2A* deletion, suggesting a higher-grade molecular profil despite its grade 2 classification. Cell proliferation was monitored for 10 days, LGG85 and LGG349 doubled every 6 days, LGG275 every 9 days, and LGG336 in more than 10 days. In comparison, three previously GB-derived cell lines[16] proliferated significantly faster, with doubling times of approximately 2-3 days (**Fig.1B**).

**Figure 1:**
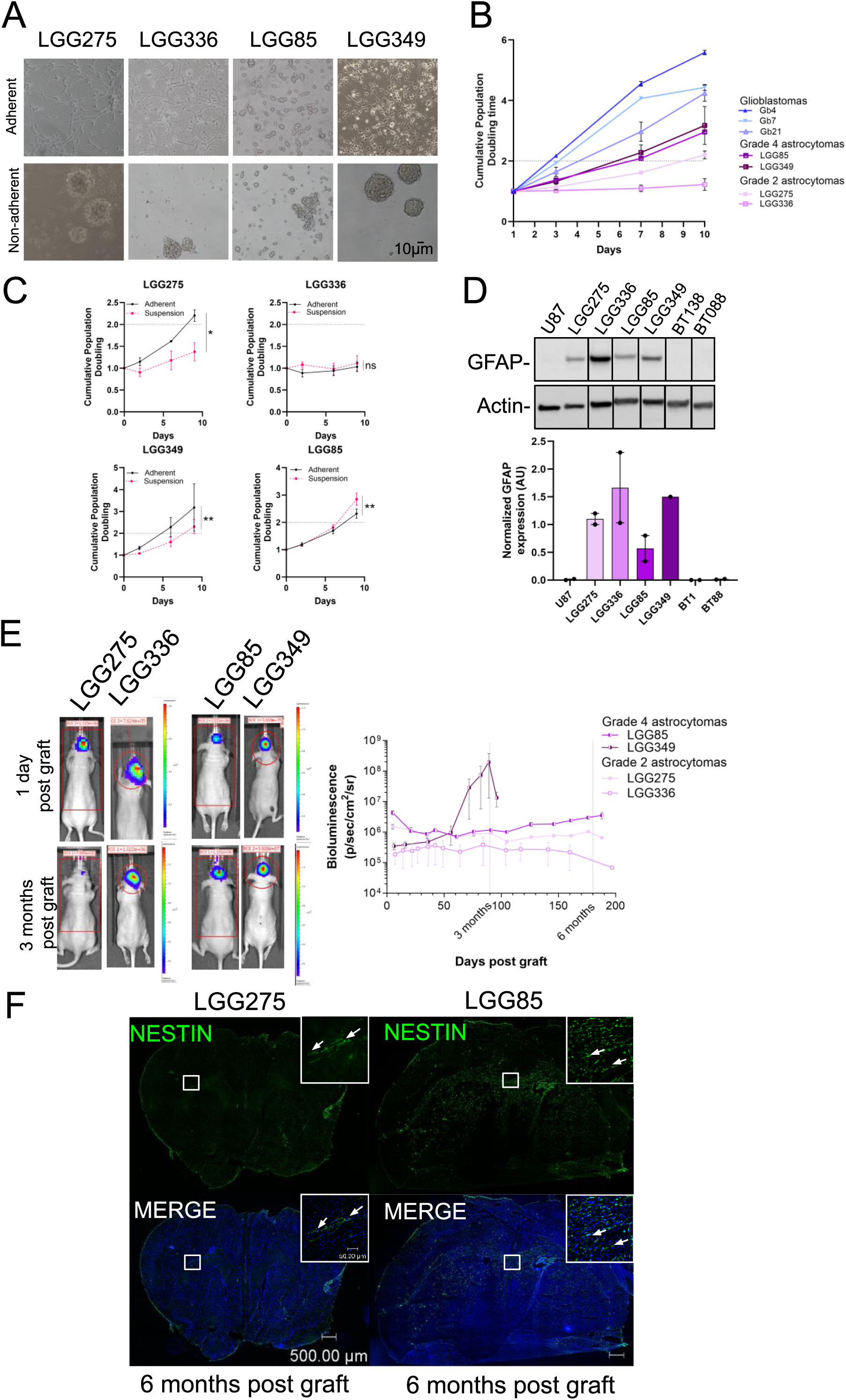
Grade 4 cell lines are more proliferative and tumorigenic than grade 2 cell lines. (A) Cell morphology of the 4 astrocytomas cell lines, in adherent on non-adherent (suspension) conditions (B) Cell doubling time of astrocytoma (n=3) and glioblastoma (n=1) cell lines in adherent conditions(C) Comparison of cell doubling time in adherent or non-adherent conditions (n =3) with GF. Each point represents mean ± S.E.M (ns = non significant, * p<0,05 et **p<0,01), as determined with a Student t-test (D) Expression and quantification (arbitrary unit) by western blot of GFAP in astrocytoma cell lines with growth factors (E) Bioluminescence visualization of astrocytoma cell lines grafted in mice (n=7) (F) NESTIN expression in astrocytoma cells (LGG275 and LGG85) in mice brain slice 6 months post-graft. Nuclei are stain with Hoechst 33342. Scale bars: 500 µm, and 50 µm for insights.

Given that glioma cell lines are frequently maintained as neurospheres under non-adherent conditions[17–19], we evaluated this alternative culture format. LGG275, LGG336, and LGG349 formed compact neurospheres, whereas LGG85 generated loose cellular aggregates (**Fig.1A**). Comparative growth analysis demonstrated that LGG275 and LGG349 proliferated more efficiently under adherent conditions, LGG85 favored aggregate growth, and LGG336 exhibited slow growth in both culture formats (**Fig.1C**). To ensure experimental consistency, all subsequent analyses were performed using adherent culture conditions. Western Blot (WB) expression analysis confirmed that all four lines expressed the astrocytic marker GFAP, distinguishing them from two reference oligodendroglioma cell lines (BT138, BT088)[17], supporting their astrocytic identity (**Fig.1D**).

We next assessed the tumorigenic potential of these lines by intracranial grafting with luciferase bioluminescence monitoring. LGG275 and LGG336 showed no detectable tumor growth during six months of observation (**Fig.1E**). Histological examination of LGG275 grafts using hNESTIN staining showed cell survival in the brain without obvious proliferative expansion. In contrast, LGG85 grafts demonstrated proliferation (**Fig.1E**) and brain parenchymal infiltration (**Fig.1F**), while LGG349 cells exhibited the highest aggressiveness, generating a strong bioluminescent signal within 3 months (**Fig.1E**).

To comprehensively characterize these lines at the molecular level, we performed whole-exome sequencing (WES) and WB analyses[20]. WES revealed that LGG85 harbored the highest mutational burden (4,172 mutations, predominantly missense variants), consistent with its prior temozolomide exposure, which is known to promote hypermutated phenotypes in gliomas (**Fig.2A**)[21]. Both WES and WB confirmed IDH1-R132H mutations in three lines (LGG275, LGG336, and LGG85), while LGG349 had lost it; a phenomenon previously documented during line establishment from *IDH1*-mutant gliomas (**Fig.2B.C**)[17]. Functional validation confirmed 2-hydroxyglutarate (2-HG) secretion in the three *IDH1*-mutant lines but not in LGG349, corroborating the genomic findings (**Fig.2D**). As anticipated for astrocytic tumors, *TP53* mutations were present across all four lines, while *ATRX* alterations occurred in three lines (**Fig.2C**). ATRX protein expression was absent in LGG275, LGG336, and LGG349 but preserved in LGG85, as U87 cell lines, used as positive control (**Fig.2B**). Notably, both LGG275 and LGG336 carried mutations in *SMARCA4*, encoding a chromatin remodeling complex component (**Fig.2C**)[6].

**Figure 2:**
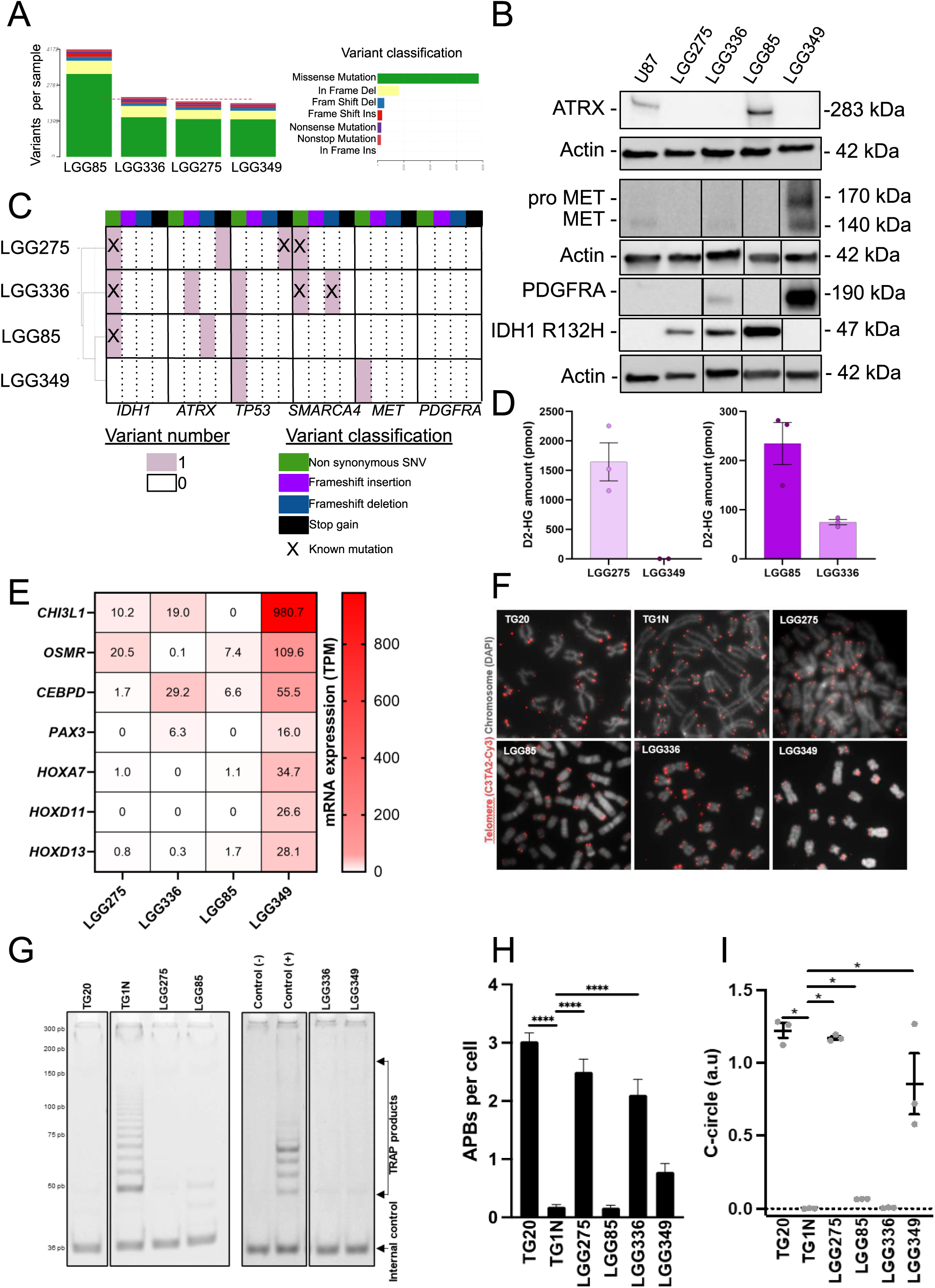
Molecular characteristics of astrocytoma cell lines. (A) Number of genetic variants in astrocytoma cell lines (B) Expression of ATRX, IDH1 R132H, proMET and MET and PDGFRA and in astrocytoma cell lines by western blot (C) Number and classification of genetic variants in *ATRX, IDH1, MET, PDGFRA* and *TP53* genes in astrocytoma cell lines after WES using VarDecrypt (D) Intracellular production of 2-HG in astrocytoma cell lines (n=3). (E) Heatmap of mRNA expression (RNA seq) of mesenchymal (*CHI3L1, OSMR, CEBPD*) and neurodevelopmental (*HOX, PAX)* genes in cell lines, n=1 (F) Representative chromosomes hybridized with a telomeric PNA probe (red) in Telo-FISH experiments. Chromosomes were counterstained with DAPI (grey) (G) Telomerase activity determined by TRAP and PAGE (H) ALT-associated PML bodies (APBs) per cell. Bars show the mean ± S.E.M (**** p<0.001, one-way-ANOVA followed by Dunnett’s multiple comparison test). (I) Detection of C-circles by C-circle assay. Bars show means ± S.E.M, with dots representing individual experimental values (n = 3) (* p<0.05, Mann-Whitney test). Telomerase-positive cells (TG1N) and ALT-positive cells (TG20) were used as controls in all experiments.

Consistent with its aggressive *in vitro* and *in vivo* behavior, LGG349 displayed molecular features characteristic of high-grade astrocytoma, including *MET* mutations and overexpression of both MET and PDGFRA (**Fig.2B.C, Fig.S1A.B**). Importantly, high MET expression was already detectable by immunofluorescence (IF) in the original patient’s tumor, confirming this alteration preceded culture adaptation (**Fig.S1C**). Consistent with its aggressive phenotype, RNA-sequencing of the four cell lines (**Fig.2E**) revealed that LGG349 expressed mesenchymal markers (*CHI3L1, OSMR, CEBPD*) alongside aberrant activation of neurodevelopmental transcription factors (*HOXA7, HOXD11, HOXD13, PAX3*). This expression signature aligns with previous reports demonstrating that neurodevelopmental/HOX programs become upregulated during malignant progression of *IDH1*-mutant gliomas[22–24].

### Evidence of alternative lengthening of telomeres in astrocytoma cell lines

*ATRX* is a crucial chromatin remodeler that maintains heterochromatin stability, especially at telomeres and repeat-rich regions. Its loss is associated with alternative lengthening of telomeres (ALT), a telomerase-independent mechanism that extends telomeres via homologous recombination. ALT occurs in about 23% of astrocytomas[25], but there are limited in vitro models for this process in these glioma subtypes[26]. To assess whether our four astrocytoma-derived cell lines display features of ALT, we examined multiple established hallmarks of this mechanism, including telomere length heterogeneity, absence of telomerase activity, the presence of ALT-associated promyelocytic leukemia (PML) nuclear bodies (APBs), and the accumulation of partially single-stranded telomeric DNA circles (c-circles) [27–30], using telomerase-positive TG1N and ALT TG20 GB lines as controls[31]. Telomere-FISH revealed heterogeneous telomere lengths, ranging from undetectable or small signals to ultra-bright foci across our four lines, in contrast to telomerase-positive TG1N cells showing homogeneous telomere foci (**Fig.2F**). TRAP assays showed that telomerase activity was undetectable in LGG275, LGG336, and LGG349, but low in LGG85 (**Fig.2G**). Immunofluorescence (IF) of PML protein combined with Telomere-FISH revealed ALT-associated PML nuclear bodies (APBs) in LGG275 and LGG336, but not in LGG85 or LGG349 (**Fig.2H**). Moreover, telomeric c-circles were detected in LGG275, LGG85, and LGG349 (**Fig.2I**). Taken together, these findings indicate that only LGG275 consistently exhibits all hallmark features of ALT, whereas the other lines display partial features, suggesting heterogeneous telomere maintenance mechanisms or potentially transient or incomplete ALT activation.

### Astrocytoma cell lines retain astrocyte-like, oligodendrocyte-like, and NPC-like cell states

Astrocytomas are heterogeneous tumors composed of distinct cellular states, including astrocyte-like and oligodendrocyte-like populations [7–9]. These tumors also contain a subset of proliferative cells resembling NPC[9]. However, the persistence of these three cell types in our glioma lines was uncertain. To address this question, we performed single-cell RNA sequencing (scRNA-sequencing) on the four glioma lines cultured with growth factors (GF). Annotation (**Table.S2**) using the scType algorithm confirmedthe presence of three major cellular states; astrocyte-like, oligodendrocyte-like, and NPC-like across all four lines (**Fig.3A.B**). To confirm the maintenance/annotation of these three subpopulations, we performed Gene Set Enrichment Analysis (GSEA). The three main identified clusters revealed molecular signatures matching astrocyte-like, oligodendrocyte-like, and NPC-like states (**Fig.S2A, Table.S3**). Typical markers for proliferation/NPC, astrocytes (e.g., *APOE, AQP4, CD44, GABBR2, ID1*), and oligodendrocytes (e.g., *DLL3, OLIG1, OLIG2, OPCML*) were predominantly expressed in the corresponding clusters. Genes defining NPC-like identity (e.g., *AURKB, E2F7*) are absent from astrocyte- and oligodendrocyte-like states (**Fig.3C.D**). Additionally, “unknown” cells were detected in all lines, with higher proportions in LGG336 and LGG349, suggesting subclonal heterogeneity (**Fig.3A**).

**Figure 3:**
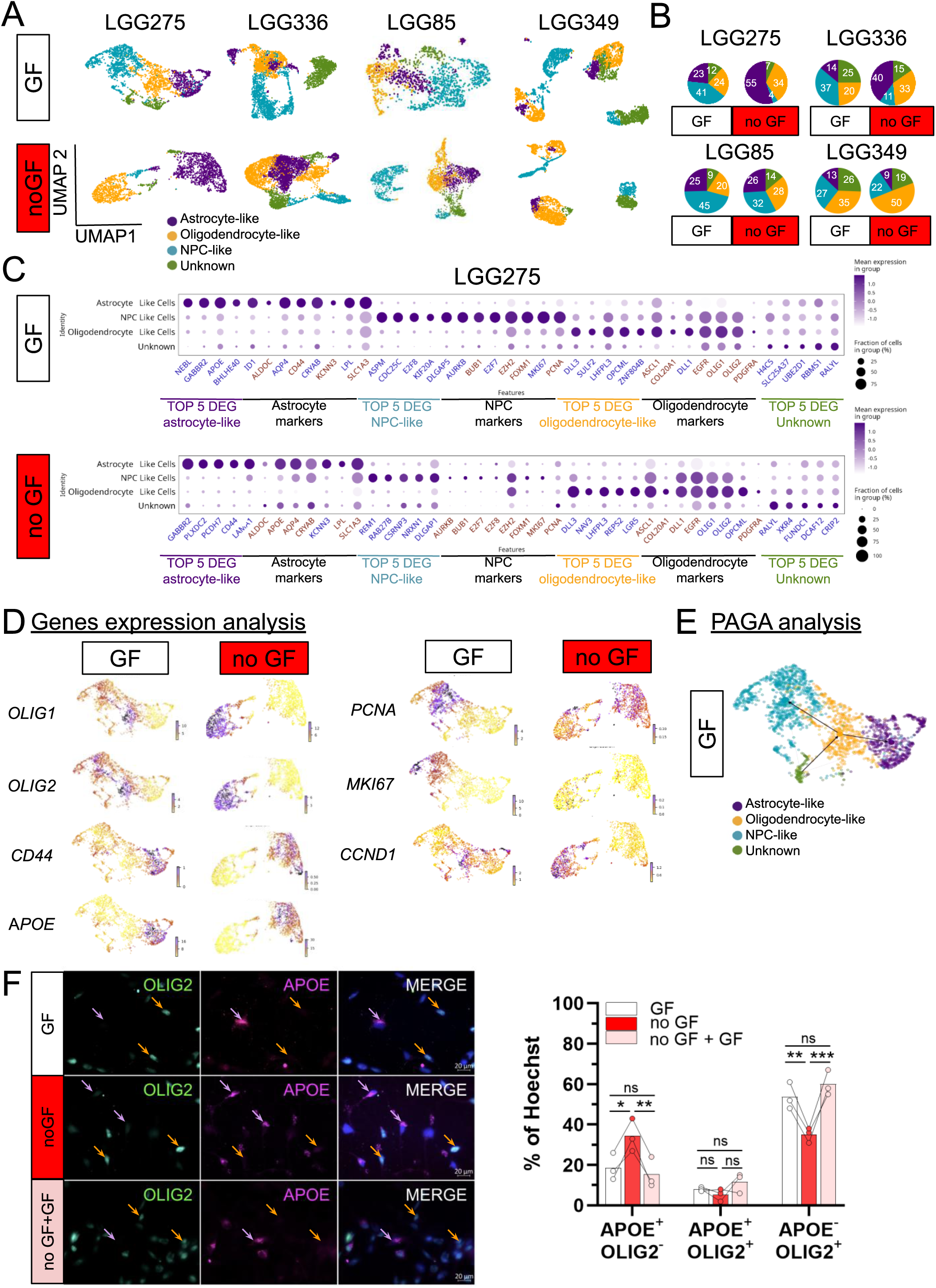
Astrocytoma cell lines exhibit multiple cellular states, reflecting the heterogeneity observed in patient tumors. (A) UMAP visualization of scRNA-seq transcriptomic clusters from four astrocytoma cell lines cultured with (GF) or without (no GF) growth factors. (B) Cellular state proportions in astrocytoma cell lines with or without GF (C) DotPlot of curated differentially expressed genes (DEG) (TOP5 + markers of astrocyte, oligodendrocyte or NPC) in each cellular states in LGG275 cell line with or without growth factors. Genes in blue are among the TOP20 DEG (D) Examples of oligodendrocyte-like (*OLIG1, OLIG2*) and astrocyte-like specific (*APOE, CD44*) gene expression measured by scRNA-seq in distinct cell subpopulations of the LGG275 cell line cultured with or without GF. Genes are differentially expressed between each subpopulation with a p.value <0.01 as determined by a Wilcoxon test (E) Partition-based graph abstraction (PAGA) analysis integrating RNA velocity to infer trajectories between cell subpopulations (F) Immunofluorescence (IF) analysis of APOE and OLIG2 expression in the LGG275 cell line, identifying “astrocyte-like” APOE+OLIG2− cells (purple arrows) and “oligodendrocyte/NPC-like” APOĒOLIG2+ cells (orange arrows) under different growth factor conditions: with (GF) or without GF (no GF) for 5 days, or without GF for 5 days followed by GF re-exposure for 5 days (no GF + GF). Right panel: Barplot show quantification as % of Hoechst+ cells (bars = mean, dots = n = 3) (*p < 0.05, **p < 0.01, \*\**p < 0.001,* multiple comparisons Tukey test).

Proportions of significant DEG-matching astrocyte-, oligodendrocyte-, or NPC-lineage reference gene lists were comparable among LGG275, LGG336, and LGG349, but decreased significantly in LGG85 with GF (**Fig.S2B, Table.S4**). Compared to LGG275, cell-type boundaries in other lines were less distinct. For example, in LGG336 and LGG349, NPC-associated genes (*EZH2, FOXM1*) were strongly found in astrocyte- and oligodendrocyte-like clusters, while oligodendroglial markers (*OLIG1, OLIG2*) were present in astrocyte-like clusters. Astrocytic markers (*CD44, CRYAB, GABBR2, KCNN3*) were enriched in astrocyte-like clusters, but extended to others (**Fig.S2C.E**). LGG85 expressed them minimally (**Fig.S2D**).

We also examined cellular states proportion after five days of GF withdrawal (noGF) which leads to decrease of proliferation in the 4 cell lines as shown by RNA sequencing and GSEA (**Fig.S3A.B**), However, While the proportions of astrocyte-like, oligodendrocyte-like, and NPC-like states were similar across the four lines cultured with GF, the situation changed markedly upon withdrawal. The NPC-like population dropped sharply by ∼10-fold (41% → 4%) in LGG275 and ∼3.3-fold (37% →11%) in LGG336, whereas the decline was much less pronounced in the two high-grade astrocytoma-derived lines, with only a ∼1.4-fold reduction in LGG85 (45% → 32%) and a ∼1.2-fold reduction in LGG349 (27% → 22%) (**Fig.2B**).This suggests that as astrocytomas progress toward higher malignancy, NPC-like cells become less dependent on exogenous growth factors for their maintenance.

To pursue downstream analyses, we prioritized LGG275 for further analysis due to its scRNA-sequencing profile resolving three distinct clusters similar to patient samples, as indicated by marker expression (*APOE, CD44, OLIG1, OLIG2, MKI67, PCNA, CCND1*) (**Fig.3C.D**)[7]. Additionally, LGG275 exhibits a significant decrease in NPC-like cells following GF withdrawal and is non-tumorigenic *in vivo*. Genetically, LGG275 has canonical changes of IDH-mutant low-grade astrocytomas: *ATRX* loss, *TP53* mutation, and IDH1-R132H substitution. Collectively, these features indicate that LGG275 resembles slow-growing astrocytomas and, in our view, provides a particularly relevant model for in-depth investigation.

To delineate relationships between cell states in LGG275, we computed a PAGA graph and oriented edges using RNA-velocity estimates from scVelo software[32]. The graph reveals a branched topology with an oligodendrocyte-like hub that is strongly connected and shows directed transitions from the oligodendrocyte-like state toward NPC-like and astrocyte-like populations (**Fig.3E**). A small “unknown” cluster shows velocity flowing into an oligodendrocyte-like node, suggesting a transient state. This indicates high plasticity, making the oligodendrocyte-like state an intermediary state between NPC-like and astrocyte-like cells.

To confirm the presence of astrocyte-like and oligodendrocyte-/NPC-like subpopulations in the LGG275 line, we used IF with APOE as an astrocyte marker and OLIG2 as an oligodendrocyte/NPC marker (**Fig.3F**). With GF, 20% of cells are APOE^+^OLIG2^-^, 57% are APOE^-^OLIG2^+^, and ∼10% co-expressed both, suggesting a transitional state between these phenotypes. Consistent with the scRNA-sequencing data, growth factor withdrawal increased the proportion of astrocyte-like (APOE+OLIG2^-^) cells to 38% and reduced oligodendrocyte/NPC-like (APOE^-^OLIG2+) cells to 39%. To explore the plasticity of these states, we reintroduced GF (noGF + GF) and found that the proportions of APOE^+^OLIG2^-^ and APOE^-^OLIG2^+^ cells returned to baseline, suggesting reversibility and dynamicity in these state transitions. Similar IF analyses using additional astrocytic markers (GFAP, FABP7) and oligodendrocyte/NPC associated markers (MASH1, OLIG1) confirmed the coexistence of astrocyte-like and oligodendrocyte-like populations in LGG275 cells (**Fig.S3C**). Flow cytometry with CD44 (astrocyte-like) and OLIG1 (oligodendrocyte/NPC-like) further corroborated these findings, revealing two distinct populations present in approximately equal proportions without GF (**Fig.S3D**). The OLIG1 and CD44 flow cytometry analysis of the other three lines did not show clear clusters (**Fig.S3D** reinforcing the idea that LGG275 better captures the cellular state heterogeneity observed in patients with slow-growing astrocytomas.

### Astrocyte-like cells define a quiescent state in LGG275 line

It is established that tumors contain subpopulations of cells that are either in a quiescent state or actively proliferating, and this also applies to gliomas[10]. However, the existence of such subpopulations in astrocytoma cell lines has not been documented. To first access cell cycle phase of LGG275 cells, we performed a cell cycle analysis using propidium iodide (PI) staining, which revealed that the vast majority of cells were not actively cycling and are in G0/G1 phase (**Fig.S4A**). To determine whether most LGG275 cells eventually re-enter the cell cycle and actively divide, we carried out dye dilution assays with or without GF. As shown in **Figure.4A**, after 20 days, cells cultured with GF had largely lost their dye fluorescence, whereas those deprived of GF retained it. This indicates that the majority of LGG275 cells entered the cell cycle over this period and that quiescent cells can be driven back into proliferation by GF, supporting the existence of a reversible quiescent state. To visually distinguish between proliferating and quiescent subpopulations in LGG275, we used IF for *CDKN1B*/P27, an indicator of quiescence, and KI67. **Figure.4B** shows two distinct populations of KI67^-^P27^+^ and KI67^+^P27^-^ cells. As expected, GF withdrawal increased the proportion of KI67^-^P27+ (quiescent) cells, while reintroduce GF restored the initial proportions, confirming that quiescent state is reversible.

To isolate quiescent cells, we engineered LGG275 cells to express a mVenus/P27 fusion protein via viral transduction[33]. Since P27 is rapidly degraded upon cell-cycle entry, fluorescence intensity enabled discrimination between quiescent (mVenus/P27 +) and cycling (mVenus/P27 −) cells. GSEA of RNA sequencing profiles from mVenus/P27 ^+^ and P27/Venus^-^ fractions showed a marked depletion of proliferation-associated genes in the mVenus/P27 ^+^ population, confirming reporter specificity (**Fig.S4B, Table.S5.S6**). After sorting and culturing mVenus/P27 + and mVenus/P27 − cells, both populations regenerated similar proportions of each type within 4 days, confirming that quiescence in LGG275 cells is reversible and dynamically regulated (**Fig.S4C**).

As our scRNA-sequencing analysis revealed several clusters, we next sought to examine the relationship between cell phenotype and proliferative or quiescent state. To this end, we used several complementary approaches. First, we applied a new classifier (**ccAFv2**)[34] providing improved resolution of G0-phase cells compared with Seurat’s *CellCycleScoring* function, which cannot reliably distinguish G0 from G1. This approach identified a population of cells in a neural G0 state within the LGG275 line (**Fig.4C**). Notably, this population mostly coincided with the astrocyte-like cluster (**Fig.4C**, left). Indeed, analysis of the four cellular states in LGG275 (**Fig.4C**, right, **Table.S7**) using this classifier showed that neural G0 (quiescent) gene program was strongly correlated to astrocyte-like phenotype (Pearson residual (PR) = 16.0), then oligodendrocyte-like phenotype (PR = 6.0), and negatively correlated to NPC-like identity (PR = -14.1). In contrast, proliferative phases (S/G2 and G2/M) were strongly associated with NPC-like phenotype (PR = 21.6 and 12.7), as expected. G1 and M/EarlyG1 programs are linked to oligodendrocyte-like cells (PR = 4.7 and 4.1); suggesting a transient phase for this population.

**Figure 4:**
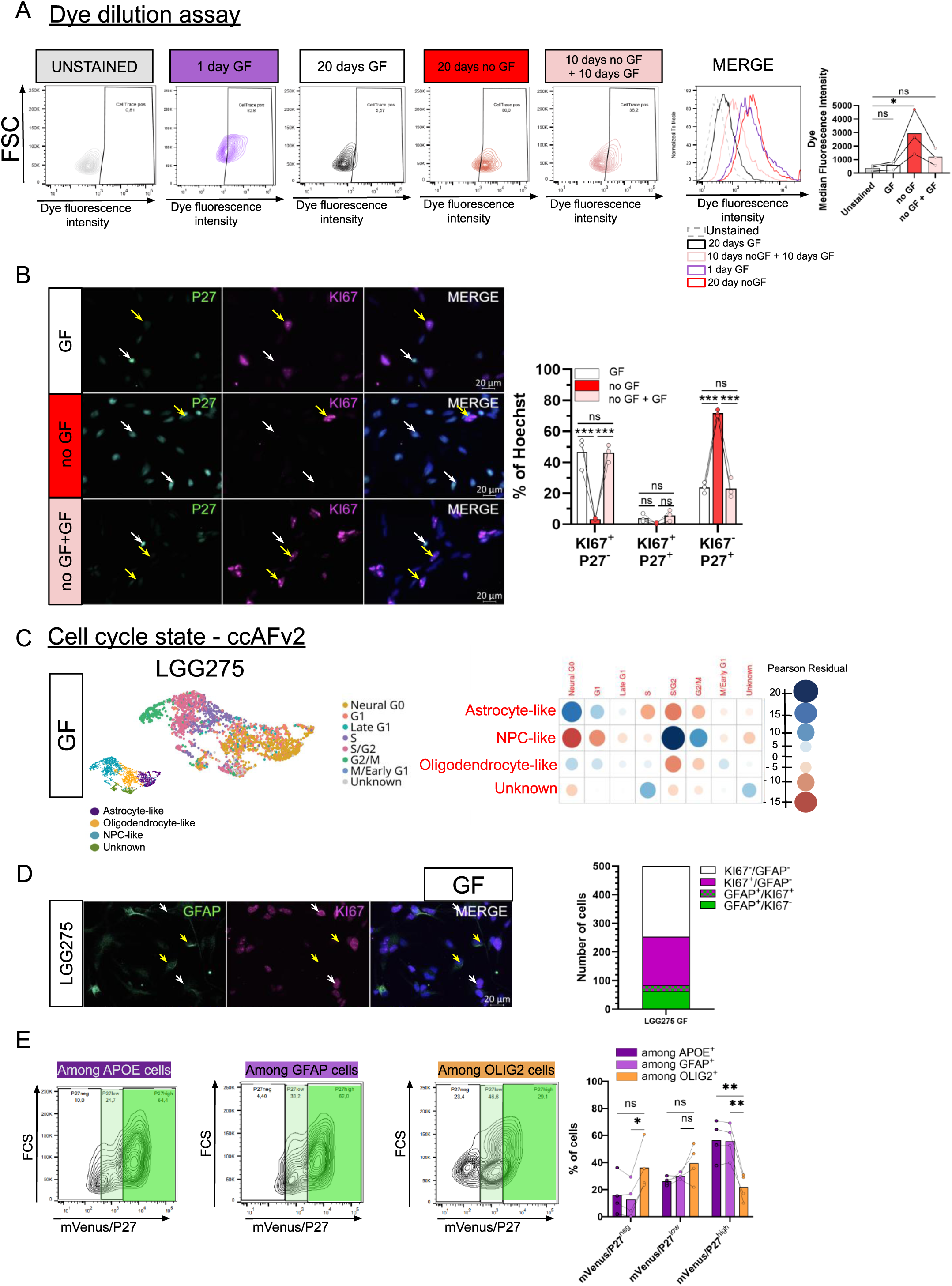
Astrocyte-like cells in the LGG275 cell line are enriched in a quiescent (Neural G0) state. (A) CellTrace dye-dilution assay by flow cytometry in LGG275 cell lines after 20 days in “GF”, “no GF” and “no GF + GF” conditions. Left: Representative flow cytometry plots (FSC vs CellTrace fluorescence) for unstained cells and CellTrace-labeled cells after 1 day GF, 20 days GF, 20 days no GF, or 10 days no GF + 10 days GF. Middle: Overlaid bars (MERGE) show CellTrace fluorescence intensity. Right, CellTrace median fluorescence intensity (MFI) quantification. (mean ± S.E.M; dots = independent experiments, n = 3); (*** p<0,001) as determined by a Mann–Whitney multiple comparisons. (B) KI67/P27 IF in LGG275 glioma cells cultured in GF, no GF, or no GF → GF conditions. Proliferating KI67+P27− cells (white arrows) and quiescent KI67−P27+ cells (yellow arrows) are indicated; nuclei (Hoechst). Right, Barplot show quantification as % of Hoechst+ cells (bars = mean; dots = n = 3); (*** p<0,001) as determined by one-way ANOVA with Tukey’s multiple comparisons. (C) Single-cell RNA-seq analysis of LGG275 cells cultured in GF. Left, UMAP colored by ccAFv2 cell-cycle state classifier, including Neural G0. Right, dot plot showing the association between phenotypic states (astrocyte-like, NPC-like, oligodendrocyte-like, unknown) and ccAFv2 cell-cycle classes (Pearson residual). Significance was assessed by a χ² test (p < 0.01). (D) IF for GFAP and KI67 in LGG275 cells grown with GF showing two non-overlapping astrocyte-like (GFAP^+^, yellow arrows) and mitotic (KI67^+^, white arrow) populations (n=1) (E) Flow cytometry analysis of the mVenus–P27 fusion reporter in LGG275 cells. Cells cultured with growth factors (GF) were stained for APOE, GFAP, and OLIG2 and analyzed by FACS to assess reporter expression. Representative density plots show mVenus/P27 fluorescence among APOE+, GFAP+, and OLIG2+ gated cells. Right: Barplot show quantification of the fraction of APOE+, GFAP+, and OLIG2+ cells across mVenus/P27 expression levels (bars = mean; dots = n = 3) (ns, not significant; *p<0.05; **p<0.01, Tukey’s multiple comparisons test).

We experimentally validated these predictions using IF and flow cytometry. Double labeling for GFAP and KI67, to identify astrocyte-like and proliferating cells, respectively, revealed that the two populations were largely mutually exclusive (**Fig.4D**). Triple KI67/APOE/OLIG2 stainings revealed that KI67 expression was predominantly associated with APOE^-^OLIG2+ cells, rather than APOE+OLIG2^--^ cells, with or without GF (**Fig.S5A**). Consistently, flow cytometry analysis of the P27/mVenus reporter line showed that P27^high^ cells were more frequently APOE+ and GFAP^+^ than OLIG2+ (**Fig.4E**), further supporting that astrocyte-like cells preferentially adopt a more quiescent state.

The relationship between cellular identity and quiescent state was also studied in other lines. The **ccAFv2** classifier revealed that cells in the neural G0 phase of LGG336 and LGG349 exhibited mixed astrocyte-like and oligodendrocyte-like signatures (**Fig.S5.B**). In the high-grade LGG85 line, which contains only a few astrocyte-like cells, neural G0 cells were instead enriched with oligodendrocyte-like markers. Thus, although the classifier consistently identified quiescent populations across all lines, their lineage identity was more heterogeneous than in LGG275. Nevertheless, GFAP and KI67 IF analyses showed reduced proliferative activity in GFAP^+^ cells, suggesting astrocyte-like characteristics in the quiescent compartment in all lines (**Fig.S5.C**).

### Astrocyte-like and oligodendrocyte-like cells in LGG275 reflect quiescent and activated neural stem-like states

The clear coexistence of astrocyte-like and oligodendrocyte-like cells within the LGG275 line provides a unique model to investigate their respective properties and interactions. ScRNA-sequencing identified CD44 and GLAST as markers for astrocyte-like cells (**Fig.3.C.D**), and used them to isolate CD44+GLAST+ and CD44−GLAST− populations, after GF withdrawal. CD44+GLAST+ cells exhibited larger, complex morphologies, while CD44−GLAST− cells were smaller and bipolar (**Fig.5A.B**). To deeply characterize these 2 populations, we first performed RNA-sequencing on each sorted subpopulation, identifying 2,431 differentially expressed genes (DEG); 1 713 enriched in CD44+GLAST+ and 718 in CD44−GLAST− cells (**Fig.5C. Table.S8**). Extending this analysis at the protein level, quantitative proteomic analysis revealed 1 324 proteins enriched in CD44+GLAST+, and 1 360 proteins enriched in CD44−GLAST− cells (**Fig.5D**, **Table.S10**). As expected, astrocytic markers (APOE, CRYAB, FABP7, GFAP, SPARC, VIM) were upregulated in CD44+GLAST+ cells, while oligodendrocytic markers (ASCL1, DLL3, EGFR, GPR17, OLIG1/2, PDGFRA, SOX4/8) dominated in CD44−GLAST− cells (**Fig.5D.E, Table.S8**). RT-qPCR confirmed their selective expression in both populations (**Fig.S6A**). GSEA showed that CD44+GLAST+ cells were enriched in astrocyte[35] and astrocyte-like[36] gene signatures, whereas CD44−GLAST− cells exhibited enrichment in oligodendrocyte[37], OPC[38], and NPC-like[36] programs (**Fig.5F, Fig.S6B**). Cytoscape network visualization of GSEA results of RNA sequencing, and Fisher associated 1D enrichment analysis of the proteome (**Fig.5G**, **Fig.S7A**, **Table.S9.S11**) revealed that astrocyte-like (CD44+GLAST+) cells were enriched in pathways related to cell communication (tight junctions, synaptic transmission, interferon response, myelination), metabolism (lipids, amino acids, fatty acids, cholesterol, glycolysis), and signaling (GPCR and calcium). Developmental pathways, including astrogenesis, were also prominent. These cells further expressed genes linked to EMT/PMT, ECM interaction, and migration, consistent with a partial mesenchymal phenotype. These metabolic and signaling features resemble those of normal astrocytes[39–44] and have been similarly reported in quiescent glioma stem cells (qGSC) from high-grade gliomas[45]. CD44−GLAST− cells have higher expression of genes related to cell cycle, stem cells, telomere maintaining, and epigenetic control, suggesting a more proliferative, NPC-like state.

**Figure 5:**
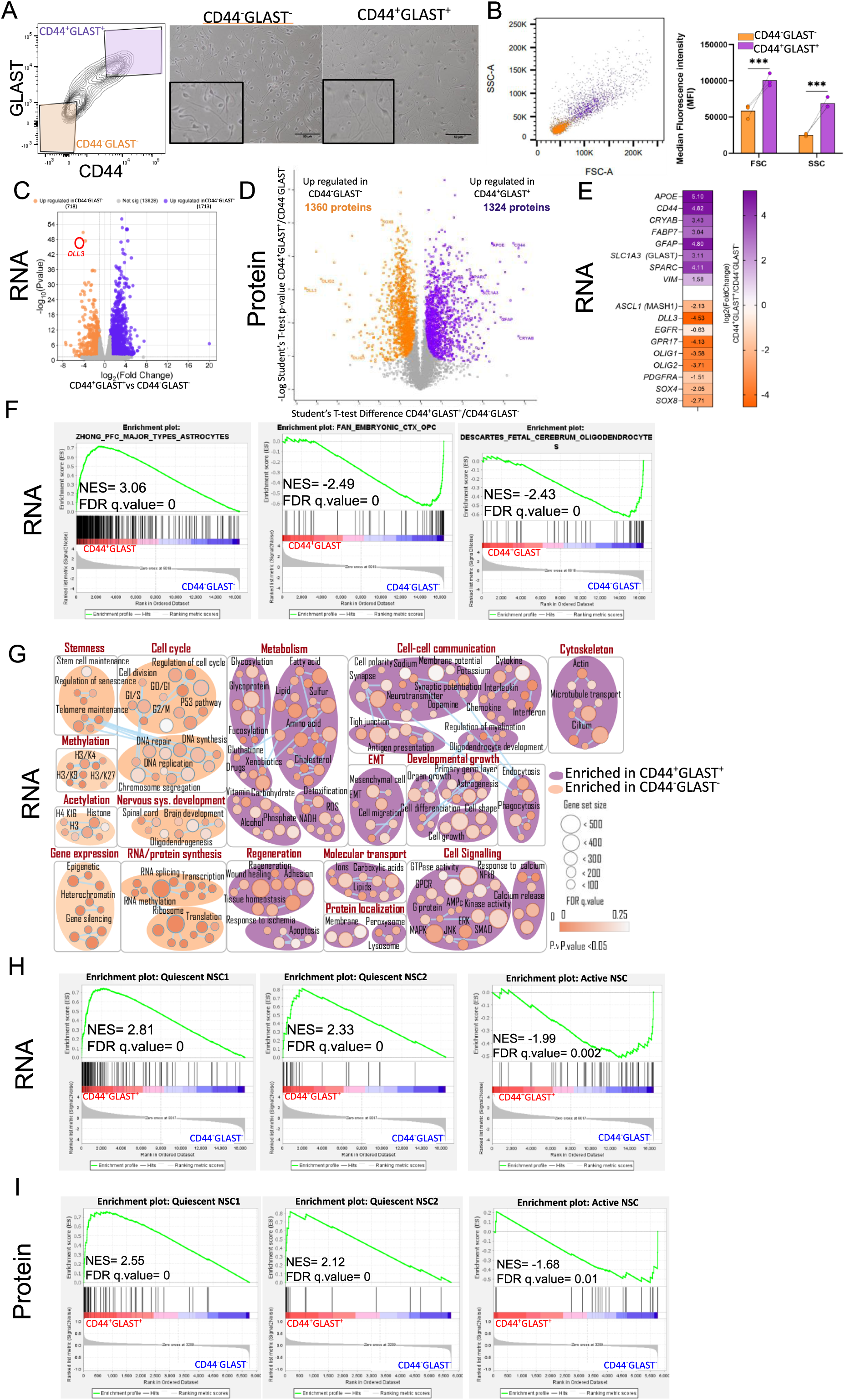
Characterisation of astro-like and oligo-like populations in LGG275 cells. (A) Morphology of astrocyte-like and oligodendrocyte-like states. Left: representative flow cytometry plot showing the gates used to isolate CD44−GLAST− and CD44+GLAST**⁺**cells. Right, phase-contrast images of the corresponding sorted populations; boxed areas indicate higher-magnification views. Scale bar, 50µm (B) Size and granularity of CD44^+^GLAST^+^ and CD44^-^GLAST^-^ cells analyzed by flow cytometry (Historams = means; dots = n = 3) (*** p < 0.001, Student’s t-test) (C) Volcano plot of differentially expressed genes between CD44^+^GLAST^+^ and CD44^-^GLAST^-^ populations (RNA sequencing). The orange and purple dots represent differentially expressed genes with a Log2(FoldChange) >1 and p.value <0.05 as determined by a Student’s t-test. (D) Volcano plot of differentially expressed proteins between CD44^+^GLAST^+^ and CD44^-^GLAST^-^ populations measured by mass spectrometry. Orange and purple dots represent differentially expressed proteins with a FoldChange>1 and p-value<0.05 as determined by a Student’s t-test. (E) Heatmap showing Log2(FoldChange) for selected astrocytic (*APOE, CD44, CRYAB, FABP7, GFAP, SLC1A3, SPARC,* and *VIM*) and oligodendrocyte/NPC-like (*ASCL1*, *DLL3, EGFR, GPR17, OLIG1, OLIG2, PDGFRA, SOX4* and *SOX8*) comparing CD44^+^GLAST^+^ versus CD44^-^GLAST^-^ populations (RNA sequencing, n=3). Differential expression was computed with DESeq2; all genes are significant (p <0.01). (F) GSEA of RNA-seq data comparing sorted CD44+GLAST+ and CD44−GLAST− subpopulations (n=3), showing enrichment of astrocyte and OPC/oligodendrocyte gene signatures in the CD44+GLAST+ and CD44−GLAST− sub-population, respectively (n=3). NES = Normalized Enrichment Score, FDR = False Discovery Rate. P.value has been calculated using the Benjamini-Hochberg method. (G) Cytoscape visualization of enriched signaling pathways (Gene Ontology) in CD44^+^GLAST^+^ and CD44^-^GLAST^-^ populations from GSEA analysis. P-value and q-value were calculated using the Benjamini-Hochberg method. (H) GSEA of RNA-sequencing data comparing CD44+GLAST+ and CD44+GLAST+ sub-populations (n=3) show enrichment of quiescent NSC1/NSC2 signatures in CD44−GLAST− cells, whereas the active NSC signature is enriched in CD44−GLAST− cells (I) Proteomics-based GSEA comparing CD44+GLAST+ and CD44−GLAST− populations (n=3) show enrichment of quiescent NSC1/NSC2 signatures in CD44+GLAST+ cells, whereas the active NSC signature is enriched in CD44−GLAST− cells.

An additional and intriguing observation concerns the relationship between these two populations and the two neural stem cell (NSC) states found in the mouse brain subventricular zone stem cell niche (SVZ). In this niche, NSC transition between quiescent (qNSC) and primed/activated (aNSC) states, with quiescence itself subdivided into qNSC1 and qNSC2 sub-states[46–51]. GSEA comparisons with SVZ datasets[48] using both our RNA-sequencing and proteomic data revealed that CD44+GLAST+ astrocyte-like cells closely resemble quiescent NSCs (qNSCs), whereas CD44−GLAST− oligodendrocyte-like cells align more closely with activated/primed NSCs (aNSCs) (**Fig.5H.I**, **Fig.S7B**). This parallel suggests that the coexistence of these two populations within LGG275 may reflect a reversible continuum between quiescent and activated NSC-like states.

### Astrocyte-like cells (CD44^+^GLAST^+^) are plastic and restore cellular heterogeneity

We assessed the stability and plasticity of astrocyte-like and oligodendrocyte-like states by separately culturing FACS sorted CD44+GLAST+ and CD44−GLAST− populations without GF, monitoring APOE and OLIG2 expression over 30 days. As shown in **Figure.6A**, the proportion of APOE+OLIG2− cells decreased from approximately 60% to 40%, while APOĒOLIG2+ cells increased from about 5-10% to 25-30% by day 7 in CD44+GLAST+ cells. By day 30, the proportions of the two populations became nearly equivalent. As cell division was virtually absent without GF (**Fig.3A.C– see MKI67 gene on Fig.3D**), these trajectories most likely represent an early phenotypic transition from APOE+OLIG2− to APOE^-^OLIG2+ states rather than proliferative expansion. Similar dynamics were observed with GF (**Fig.S8A**). We performed the same experiment using CD44−GLAST− cells (**Fig.6B.S8B**). One day after sorting, 80% of these cells were APOĒOLIG2+, and their phenotype remained largely stable, with only a slight increase in APOE expression over time. We confirmed these trajectories by tracking the maintenance or loss of CD44 and GLAST markers in the two purified populations. As shown in **Figure.6C**, CD44+GLAST+ cells gradually lost these markers, reconstituting a CD44−GLAST− population. In contrast, CD44−GLAST− cells were unable, under these conditions, to reacquire CD44+GLAST+ markers. These findings demonstrate that astrocyte-like cells display high phenotypic plasticity, enabling them to transition toward oligodendrocyte-like states and regenerate CD44−GLAST− populations. In contrast, oligodendrocyte-like cells are more phenotypically stable and lack the capacity to revert to the astrocyte-like state under the tested conditions.

**Figure 6:**
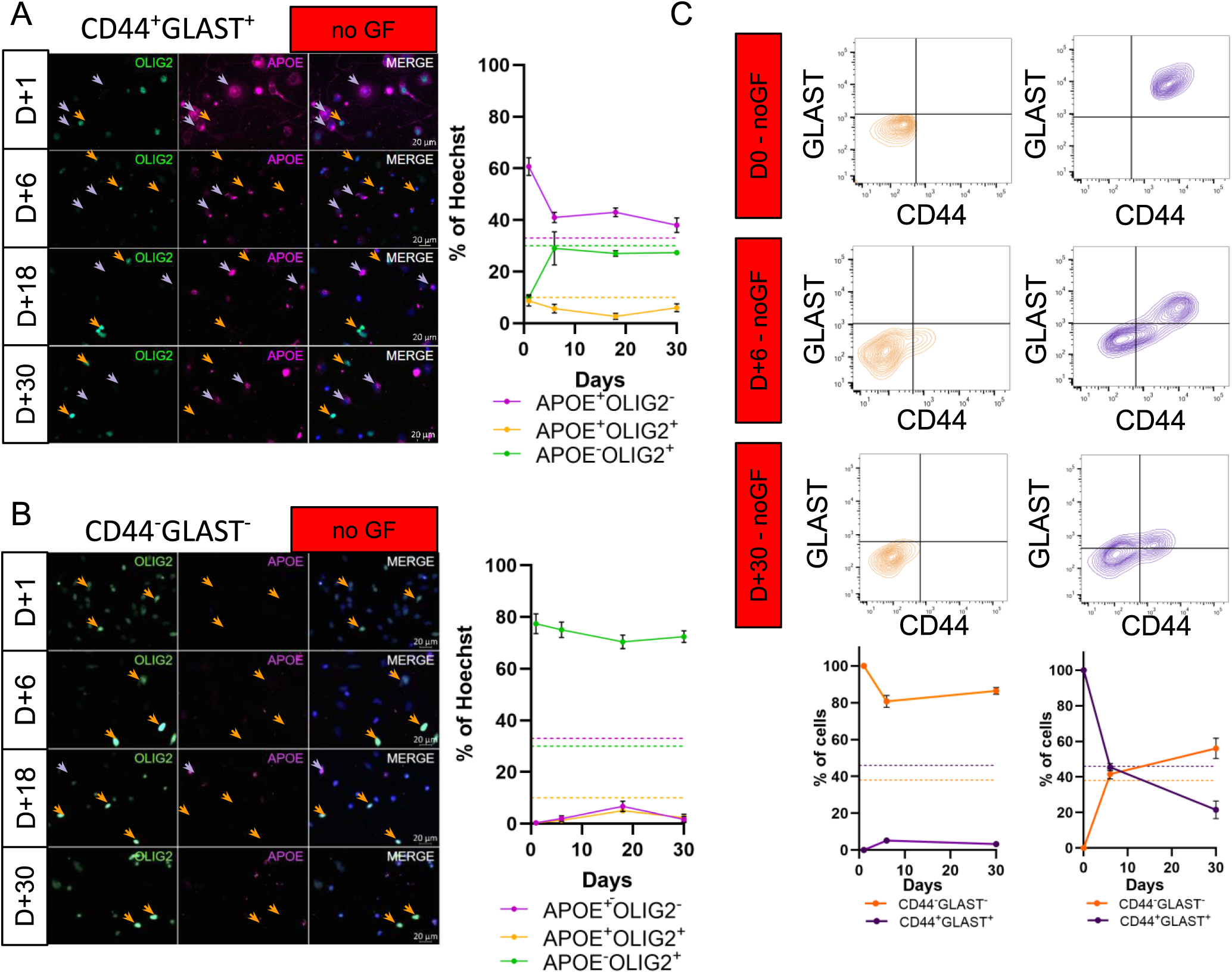
Astrocyte-like cells are plastic and play a role in the cellular heterogeneity formation in the LGG275 line. (A) IF analysis of APOE and OLIG2 expression in the LGG275 cell line, identifying “astrocyte-like” APOE+OLIG2− cells (purple arrows) and “oligodendrocyte/NPC-like” APOĒOLIG2+ cells (orange arrows) without GF, in CD44^+^GLAST^+^ and (B) CD44^-^GLAST^-^ populations, 1, 6, 18, and 30 days after cell sorting. Barplot show quantification as % of Hoechst+ cells (dots on curve = mean ± S.E.M, n=3) Flow cytometry expression of CD44 and GLAST markers in the LGG275 cell line in CD44^+^GLAST^+^ and CD44^-^GLAST^-^ populations, 6 or 30 days after cell sorting, in the absence of growth factors. Barplot show quantification as % cells (dots on curve = mean ± S.E.M, n=3)

### NOTCH-driven plasticity shapes cellular states in LGG275

The LGG275 line culture contained multiple cellular states, leading us to hypothesize that they might interact closely through direct contact or soluble factors like cytokines. In the previous experiments, we noted that oligodendrocyte-like cells (CD44−GLAST−) failed to restore cells expressing CD44, GLAST, and APOE, suggesting a lack of essential signals.

To analyze ligand/receptor interactions among LGG275 cells, we used the CellChat tool[52]. Astrocyte-like cells exhibited the highest signaling activity, mainly in an autocrine manner (124 interactions, strength 2.62), but also via paracrine signaling, sending 75 signals to oligodendrocyte-like cells (2.53) and 74 signals to NPC-like cells (1.88) (**Fig.7A**). Astrocyte-like cells therefore send more signals and engage in more interactions than oligodendrocyte-like and NPC-like populations.

**Figure 7:**
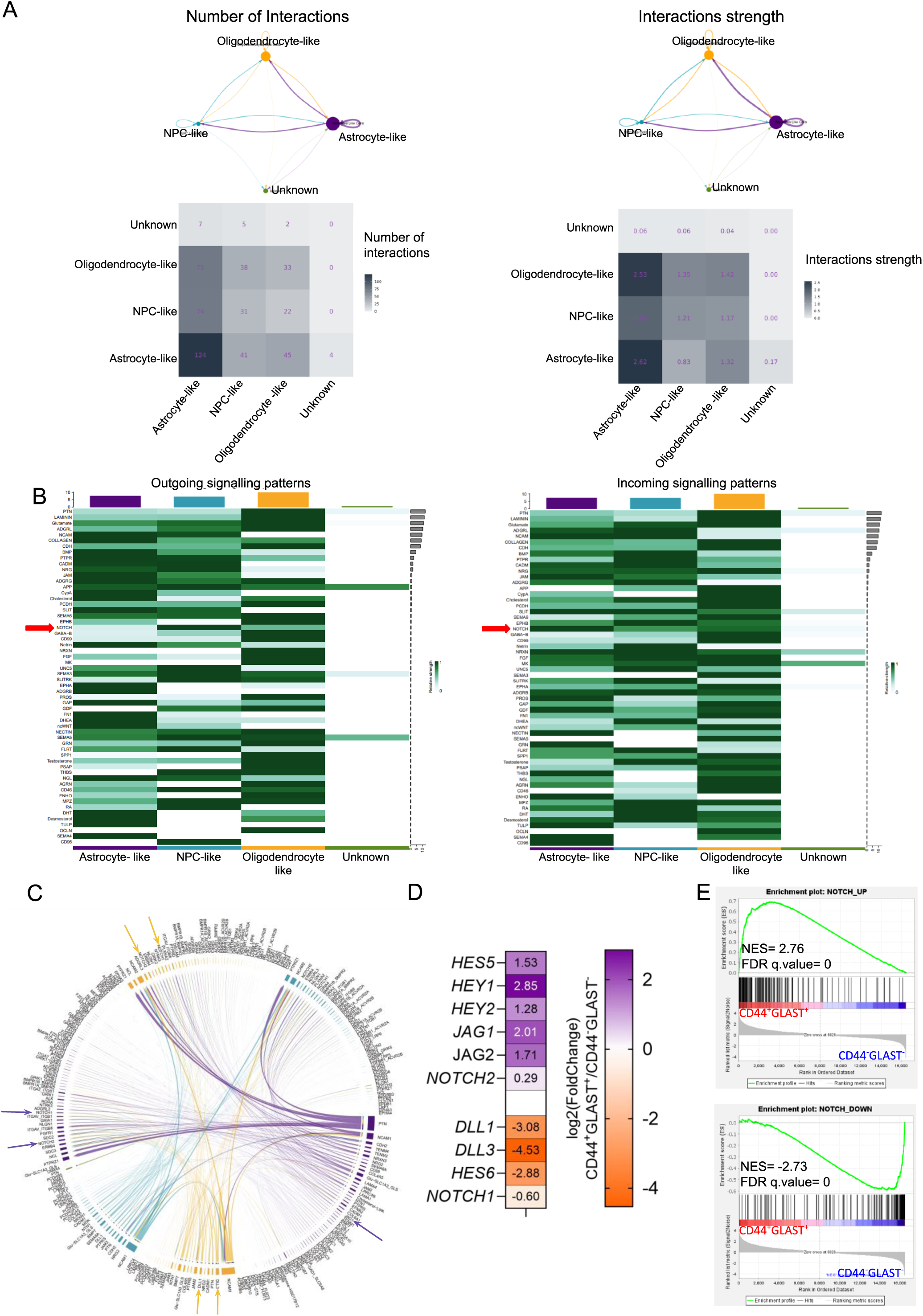
NOTCH pathway is a pathway of interaction between different cell states in the LGG275 line. (A) String diagrams and heatmaps representing the number and strength of autocrine and paracrine cell interactions in each cell subpopulations growth with growth factors. Analysis performed using the CellChat v2.1.2 tool on scRNA sequencing cell data (B) CellChat heatmap of outgoing and incoming signaling patterns inferred from scRNA-seq data. Rows indicate ligand–receptor signaling pathways; columns represent the indicated cell subpopulations. Color intensity denotes the normalized communication strength (darker = stronger). The top bar plots summarize the overall contribution of each subpopulation, and the right bar plot ranks pathways by overall information flow. Notch pathway is indicated by red arrow. (C) CellChat chord diagram showing inferred ligand–receptor communication among cell subpopulations (colors as in prior panels) from scRNA-seq data. Chord thickness reflects interaction strength, and outer labels indicate key ligand/receptor genes (arrows highlight NOTCH receptors and ligands). (D) Heatmap of canonical Notch signaling pathway genes differentially expressed between CD44+GLAST+ and CD44−GLAST− cells. Values represent log2 fold changes (CD44+GLAST+ / CD44−GLAST−) obtained from differential expression analysis performed using DESeq2 (volcano plot). All genes are significant (p <0.01). (E) GSEA analysis performed in CD44+GLAST+ and CD44−GLAST− populations (n = 3) using NOTCH_UP and NOTCH_DOWN gene sets derived from LGG275 cells following DLL4-mediated Notch activation (Table.S14) NES = Normalized Enrichment Score, FDR = False Discovery Rate.

Among the identified signaling pathways (**Fig.7B**), the NOTCH pathway stood out due to its importance in glioma cell quiescence and phenotype, as well as its role in astrocyte differentiation from NSC[7,16]. CellChat and RNA-sequencing analysis indicated that astrocyte-like cells expressed *NOTCH1/2* receptors as well as the ligand *JAG1*, while oligodendrocyte-like cells expressed NOTCH receptors alongside the ligands *DLL1* and *DLL3* (**Fig.C.D**). RNA sequencing data revealed that astrocyte-like cells had higher levels of downstream NOTCH1 transcription factors (*HEY1, HEY2, HES5*) than oligodendrocyte-like cells, which were enriched in *DLL3* and *HES6*. To further assess NOTCH pathway activity in CD44+GLAST+ and CD44−GLAST− cells, we performed GSEA on the RNA-sequencing datsa from these two populations using two custom gene sets representing genes significantly up- or downregulated following DLL4 ligand–induced NOTCH activation in LGG275 (**Table.S12**). The results shown in **Figure.7E** indicate that CD44+GLAST+ cells are enriched for genes activated by the NOTCH pathway, in contrast to CD44−GLAST− cells.

These observations prompted us to examine the plasticity of CD44+GLAST+ and CD44−GLAST− cells in response to NOTCH pathway activation and inhibition. To do this, both populations were cultured with NOTCH DLL4 ligand for six days, then APOE and OLIG2 expression were assessed by IF. As shown in **Figure.8A**, DLL4 treatment significantly increased the APOE+ OLIG2+ astrocyte-like cells from ∼39% to ∼57%, while significantly decreasing the oligodendrocyte-like/NPC-like APOĒ OLIG2+ fraction by about half. A similar shift was observed in CD44−GLAST− cells, suggesting that NOTCH1 activation strongly promotes astrocyte-like cell state (**Figure.8A right panel)**.

Conversely, **Figure.8B** shows that treatment of CD44+GLAST+ cells with two distinct NOTCH inhibitors (DAPT, LY411575) consistently led to a marked decrease in the APOE+OLIG1^-^ fraction and a robust induction of OLIG2 expression. In contrast, NOTCH inhibition had little or no effect on APOE and OLIG2 expression in CD44−GLAST− cells, suggesting a weakly active or inactive NOTCH pathway in this population.

### DLL3 regulates proliferation and phenotype of LGG275 cells

Finally, since the NOTCH pathway emerged as a regulator of astro-like vs oligo-like states within the LGG275 line, we sought to identify potential modulators of this signaling cascade. We focused on DLL3, a well-known negative regulator of NOTCH activity[53]. In proliferating cultures, scRNA-sequencing showed that DLL3^+^ cells were mainly oligodendrocyte-like, a pattern that remained without GF (**Fig.S9B**). Bulk RNA-sequencing and proteomic analysis confirm preferential DLL3 expression in oligodendrocyte-like cells (**Fig.5C.D.H**). IF shows that DLL3 is expressed in a subset (30-40%) of cells, concentrated in the Golgi, consistent with previous reports. Co-staining with OLIG2 indicates that all DLL3+ cells are OLIG2+, and this association persists without GF (**Fig.S9A**). In proliferating cultures, DLL3+ cells displayed mixed KI67 and P27 expression profiles (**Fig.S9C**), suggesting that this population may occupy an intermediate cell-cycle state.

To gain insight into the role of DLL3 in regulating LGG275 cell proliferation, we transduced the cells with a lentiviral vector overexpressing DLL3, which approximately doubled the proportion of DLL3+ cells in the culture (**Fig.8C**). This resulted in a significant increase in KI67+ cells, alongside a decrease in APOE+ OLIG2− cells and an increase in APOĒ OLIG2+ cells (**Fig.8C.D**). GSEA of RNA-sequencing data confirmed these changes, showing enrichment of NPC and proliferation signatures in cells overexpressing DLL3, while decreasing astrocyte-associated gene signatures (**Fig.8E.F, Tables.S13.S14**). Overexpression of DLL3 in a second line, LGG336, which barely expresses this gene, led to an increase in KI67^+^ cells (**Fig.S10A**) and a decrease in GFAP^+^ cells (**Fig.S10B**). Together, DLL3 emerges as a modulator of lineage plasticity and proliferation in the two cell lines we explored.

**Figure 8:**
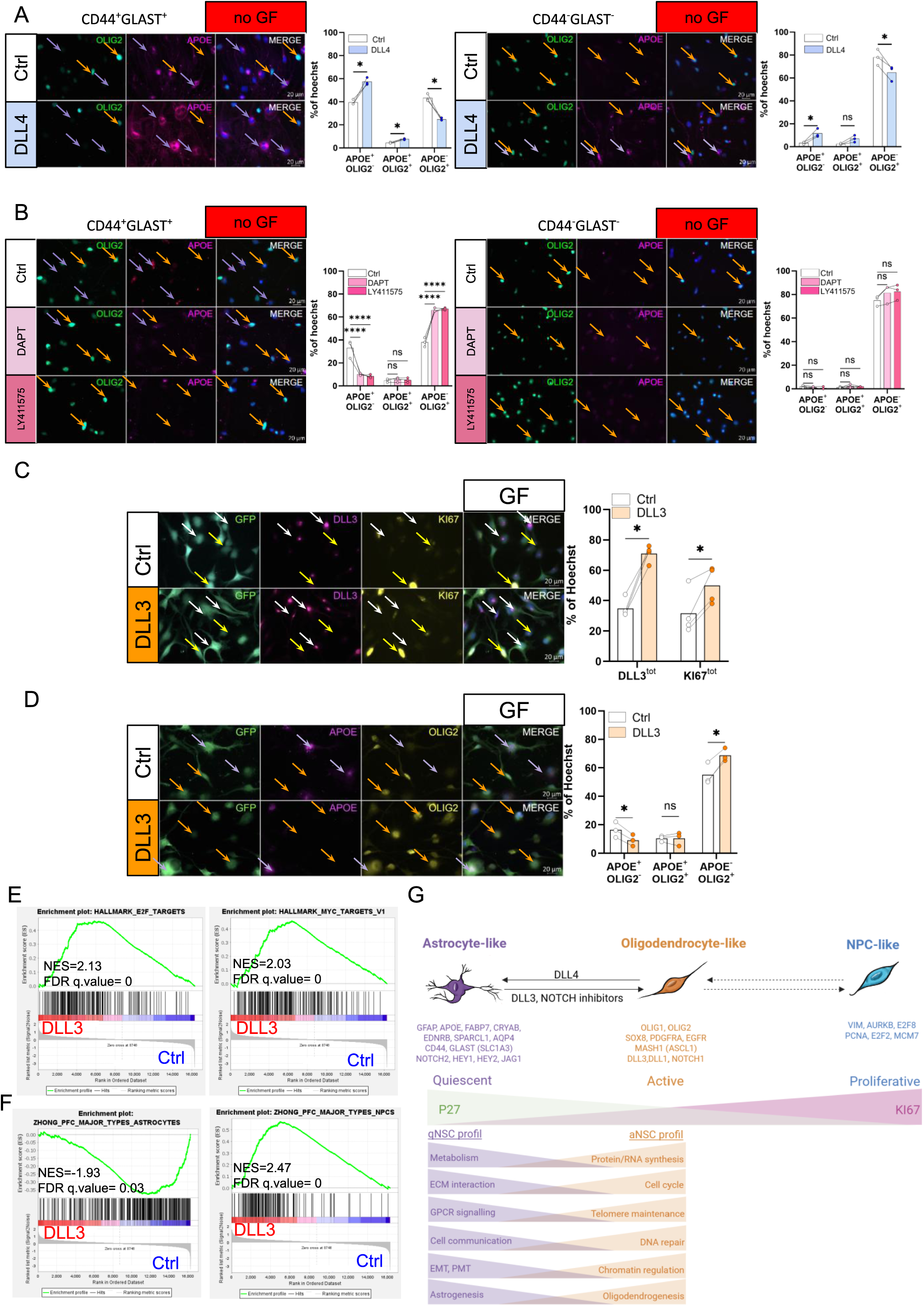
The NOTCH pathway is involved in the cellular plasticity of astrocyte-like and oligodendrocyte/NPC-like cells. (A,B) IF analysis of APOE and OLIG2 expression in the LGG275 line, identifying “astrocyte-like” APOE+OLIG2− cells (purple arrows) and “oligodendrocyte/NPC-like” APOĒOLIG2+ cells (orange arrows) without GF, in CD44^+^GLAST^+^ and CD44^-^GLAST^-^ populations. Cells were treated with the NOTCH ligand DLL4 (A) and γ-secretase inhibitors (DAPT, LY411575) (B). (C) IF analysis of DLL3 and KI67 expression in the LGG275 line identifying DLL3^+^ (white arrows) and mitotic KI67^+^ (yellow) populations with GF, after overexpression of DLL3. (D) IF analysis of APOE and OLIG2 expression in the LGG275 line, identifying “astrocyte-like” APOE+OLIG2− cells (purple arrows) and “oligodendrocyte/NPC-like” APOĒOLIG2+ cells (orange arrows) with GF, after overexpression of DLL3. For A, B, C and D, bar plots show quantification as % of Hoechst+ cells (Bars = mean, dots = n = 3) (ns p>0.05, *p<0.05, and ****p<0.0001, Dunnett’s multiple comparison test (A) or Student’s multiple test (B,C,D). GSEA analyses showing enrichment of proliferation-associated signatures (E2F and MYC targets) following DLL3 overexpression (E), and differential enrichment of astrocyte and NPC gene signatures in control (Ctrl) and DLL3-overexpressing conditions, respectively (F) (n = 3). NES = Normalized Enrichment Score; FDR = False Discovery Rate. (G) Schematic summary of the cellular states identified in the LGG275 astrocytoma cell line. Astrocyte-like cells correspond to a quiescent state that can transition to an active oligodendrocyte-like state and subsequently to a proliferative NPC-like state following DLL3 or NOTCH pathway inhibition. Conversely, activation of the NOTCH pathway promotes the maintenance or re-entry into the quiescent astrocyte-like state. These three states display molecular and functional profiles resembling quiescent (qNSC), activated (aNSC), and proliferative neural stem cells (NSC), respectively.

## Discussion

In this study, we established and comprehensively characterized four newly generated cell lines derived from IDH-mutant astrocytomas. Deep analysis of the slow-growing astrocytoma LGG275 cell line provided several key insights into the biology of IDH-mutant astrocytomas, which are summarized in **Figure.8G**.

First, glioma cell lines established under serum-free conditions are typically maintained in non-adherent cultures, allowing the formation of gliospheres[17,18]. However, we observed that for two lines (LGG275 and LGG349), this culture mode was not favorable for cell expansion. When grown on laminin, these cells displayed improved growth, suggesting that integrin-mediated signaling may be involved and could confer a proliferative advantage, as previously reported for non-tumoral human neural stem cells[54]. In contrast, the LGG336 line showed no major difference between the two culture modes, whereas LGG85 grew preferentially under non-adherent conditions. These findings highlight heterogeneity in adhesion-dependent growth requirements among the lines, potentially reflecting differences in the signaling pathways that are active in each of them.

Second, an important observation arises from the comparison between lines derived from low-grade and high-grade tumors, which respectively fail to form or successfully form tumors *in vivo*, thereby modeling the behavior of the original patient tumors. The LGG85 line exhibits a very high number of mutations, likely as a consequence of temozolomide treatment[21], whereas LGG349 shows strong expression of C-MET and PDGFRA, both known oncogenic drivers of tumor progression[6]. Moreover, LGG349 expresses developmental genes (such as HOX genes) that are normally silent in the adult brain[55]. These findings suggest that multiple distinct mechanisms can underlie tumor progression, and that these two lines will be valuable tools to dissect the molecular pathways driving malignancy. It is also important to note that although the LGG85 and LGG349 lines display high-grade characteristics, they grow more slowly than classic GB cells, underscoring the remarkable heterogeneity among high-grade gliomas, which likely results from distinct mechanisms of transformation. A salient feature distinguishing the lines derived from low- and high-grade gliomas concerns the persistence of NPC-like cells upon GF withdrawal. In both high-grade-derived lines, the proportion of NPC-like cells remains largely unchanged in the absence of GF, whereas it is markedly reduced in LGG275 and LGG336. The mechanisms sustaining NPC-like cell persistence under growth factor deprivation remain to be elucidated but are consistent with the cellular behaviors observed in patient tumors[8].

Third, scRNA-sequencing and IF analyses revealed, quite unexpectedly, the coexistence of multiple transcriptional states within each line, corresponding to astrocyte-like, oligodendrocyte-like, and NPC-like phenotypes similar to those identified in patient-derived gliomas[7–9]. The boundaries between these states were particularly sharp in the LGG275 line but became blurred in the high-grade lines, suggesting that tumor progression is associated with a loss of cell state specificity. This loss may stem from extensive epigenetic remodeling or disruptions in the underlying gene regulatory networks caused by patient-specific mutations. Using LGG275 as a reference model, we identified two surface markers, CD44 and GLAST that can be used to isolate astrocyte-like cells for functional and molecular analyses. These experiments revealed that astrocyte-like cells tend to reside in a more quiescent state but retain the capacity to exit quiescence and transition toward an oligodendrocyte-like phenotype. Morphologically, astrocyte-like and oligodendrocyte-like populations are also clearly distinct: astrocyte-like cells are larger and more flattened, whereas oligodendrocyte-like cells display a bipolar, elongated morphology. Recent study[56] has demonstrated that oligodendrocyte-like tumor cells make a predominant contribution to glioma infiltration. Their bipolar morphology observed here may therefore reflect an adaptation that facilitates migration and invasion within the brain parenchyma.

Fourth, transcriptomic and proteomic comparisons between astrocyte-like and oligodendrocyte-like populations further revealed striking parallels with SVZ NSC in their quiescent (qNSC) and activated (aNSC) states, respectively. This similarity extends, for instance, to metabolic programs: based on bioinformatic analysis of the proteomic data, astrocyte-like cells, like qNSC, preferentially rely on fatty acid oxidation; a hallmark of quiescent stem cells[46]. In the adult SVZ, the establishment and maintenance of qNSC are tightly controlled by the NOTCH signaling pathway[57–59]. In line with this, our analyses indicate that NOTCH signaling also plays a central role in regulating the balance between astrocyte-like and oligodendrocyte-like populations in LGG275 cells. Among the NOTCH-related molecules, we identified DLL3 as a particularly interesting regulator. DLL3 is known to act as an inhibitory ligand of the NOTCH pathway[60]. Consistently, it was found to be very specifically expressed by oligodendrocyte-like cells at both the RNA and protein levels, as confirmed by single-cell RNA-seq, IF, and proteomic data. Functionally, DLL3 overexpression reduced the expression of some astrocytic genes while promoting the expression of proliferation-associated genes, suggesting that DLL3 contributes to maintaining the activated, proliferative state of oligodendrocyte-like cells by dampening NOTCH activity[60]. These findings point to DLL3 as a potential marker and functional modulator of the oligodendrocyte-like, invasive glioma cell population.

### Limitations

This study has certain limitations. The lack of access to the patients’ germline and primary tumor DNA prevented a comprehensive mutational analysis. Such a comparison would have been important to distinguish true somatic mutations from genetic alterations acquired during cell line establishment. Furthermore, isolating additional cell lines resembling LGG275 would be valuable to confirm the representative nature and biological fidelity of this model for slow-growing IDH1-astrocytoma.

### Conclusion

We established and deeply characterized four IDH-mutant glioma cell lines that recapitulate molecular and phenotypic features of patient tumors, including partial or complete activation of ALT, canonical mutations (IDH1, TP53, ATRX, MET), and grade-consistent growth behaviors *in vitro* and *in vivo*. These cell lines, enriched with detailed multi-omics annotations, provide a powerful toolkit for glioma research. They enable the dissection of key pathogenic mechanisms, including ALT biology, the transition to high-grade disease, and sensitivity to emerging therapies. In our view, LGG275 stands out as a pertinent model for studying slow-growing IDH-mutant astrocytomas.

## Author Contributions

LG and JPH conceptualized the study and designed the methodology with the help of DP. LG performed all the experiments with the help of other coauthors. SH, CL, DP and KAC helped with cell culture and IF analysis. CGB performed all the ALT experiments and figures, with the help of LRG, FDB and MAM. DS performed all the single cell RNA sequencing analysis, with the help of SO and CP for the ccAFv2 algorithm. MC and LZ contributed to bioinformatics analyses and conceptual input. SH performed most of the WB analysis. KAC and CR performed the histological staining and analysis of mouse brain and patient tumor resections. LS and JK performed the 2-HG measurement assays. KE, SU and MS performed mass spectrometry and proteomics analysis. KD performed the in vivo experiment under supervision of AI and MV. HD performed all the surgeries and provided the patient tumors samples. VR provided the diagnostics and clinical information of the cases. LG and JPH wrote and corrected the manuscript, with the help of CGB for the ALT part. All authors reviewed and approved the final version of manuscript.

## Acknowledgement

We acknowledge K. Ligon, O. Abigail (Dana-Farber Institute Harvard, US) and E. Huillard, (ICM, Paris) for providing BT138 oligodendroglioma cell line. We acknowledge S. Weiss, A. (UofC, Calgary, Canada) for providing BT088 oligodendrogliomas cell line. We thank the MRI-MGX (supported from the France Génomique National Infrastructure (ANR-10-INBS09). We acknowledge the PPM, S. Urbach, M. Seveno, S. Chaumont-Dubel, K. El Koulali for mass spectrometry analyses. We thank the Montpellier Vectorology Platform (MRI-PVM, C. Lemmers) for lentivirus production, and the MRI Imaging and Flow Cytometry Platform (A. Sarrazin, M.-P. Blanchard, G. Galtier, S. Viala, M. Boyer-Clavel) for technical support.

## Material and methods

Product reference list can be found in Table.S15

Additional Material and method can be found in supplemental files, as well as products list references

### Patient-derived Cell Lines

Astrocytoma tumor biopsies (**Table.S1**) were collected from patients at CHU-Hôpital Gui de Chauliac in Montpellier, with their informed consent. Professor Hugues Duffau operated, and Professor Valérie Rigau, a neuropathologist, classified the gliomas using the WHO 2016 then 2021 guidelines[61]. The four astrocytoma cell lines: LGG275[7], LGG85[15], LGG336, and LGG349 were derived in the laboratory following the protocol of Azar et al., 2018 and Aguilar et al., 2025[14]. Tumor tissue was cut into small pieces in a PBS. The pieces were placed in wells of a 24-well plate treated with poly-D-lysine (PDL) and murine laminin. The three glioblastoma lines: Gb 4, Gb7, and Gb21 were also established in the laboratory[16]. Collaborators provided the oligodendroglioma lines BT138 and BT088. Derivation protocols are published[17,62]. We used TG1N and TG20 cell lines, which are positive for telomerase activity and alternative lengthening of telomeres, respectively, as controls. These lines were cultured as previously described [31,63].

### Cell Culture

Cells are cultured in a basal medium DMEM-F12 supplemented with N2, L-Glutamine, B27, and antibiotics. The medium is enriched with EGF, FGF_2_ and heparin for cell proliferation. A medium without these factors is also used to induce cell quiescence. Cultures are incubated at 37°C with 5% CO_2_. Cells are dissociated in Trypsin-EDTA 0.25X followed by a trypsin inhibitor, CaCl_2_, and DNAse I. Cells are then seeded on a support treated with Poly-D-Lysine and murine laminin at a density of 30 000 cells/cm^2^ for experiments.

### Cell Counting

To determine the growth rates of different cell lines, cells were cultured at a density of 20,000 cells/cm^2^, either on PDL and murine laminin, or in wells treated with polyHEMA to promote neurospheres formation. Every 1, 3, 7, and 9 days, cells were detached using 10X trypsin. Cell counts were obtained using a Z2 cell counter.

### Brain Grafts of Cell Lines

All animal procedures were performed by the Gliotex team (ICM, Paris) under procedure 17503. For orthotopic models, Astrocytoma cells were transduced with a virus containing the luciferase gene[64]. Cells were implanted (140,000 cells/2μL) into the brains of female athymic Nude mice (Reference Envigo: Hsd: Athymic Nude-Foxn1nu) at least 7 weeks old. Astereotaxic apparatus was used to inject cells into the right caudate-putamen nucleus. Luciferase intensity was monitored to analyze tumor progression. Bioluminescence acquisition was performed 10 minutes after luciferin injection using the IVIS spectrum Xenogen, and imaged were analyzed using Living Image^®^ acquisition software.

### Histological Analysis of Brain Grafts

Mice were sacrificed by cervical dislocation after gas anesthesia (Isofluorane). The brain was extracted and frozen at -80°C immediately after sacrifice to limit tissue degradation. Brain sections of 10 μm thickness were made using a cryostat. For histological analysis, brain sections were fixed with PFA 4% then dry at 4°C. Fixed tissues were permeabilized and fixation sites blocked with PBS -10% BSA, 0.3% Triton solution. To identify human astrocytoma cells, sections were incubated with a primary antibody solution (PBS 1% BSA, 0.1% triton) α-NESTIN overnight at 4° C. Sections were then incubated with a secondary antibody solution (anti-rabbit) for one hour at room temperature, and nuclei were stained with a Dapi solution. Slides were then mounted with Fluoromount-G. Sections were scanned and acquired using the Leica DM6B Thunder.

### Western Blot Analysis

Cells were scraped on ice in cold PBS then lysed using a RIPA lysis buffer, containing protease and phosphatase inhibitor cocktail. Samples were incubated on ice and centrifuged at 14,000 rpm. Protein concentrations in the supernatant were measured with a BCA protein assay. Proteins were separated by SDS-PAGE and transferred to a PVDF membrane. After blocking, membranes were incubated with the primary antibodies listed in **Table.S16** overnight at 4° C, then with peroxidase-conjugated secondary antibodies. Protein bands were revealed using the ECL kit. Images were taken using the ChemiDoc^TM^ XRS imaging system and analyzed using ImageLab software.

### Measurement of Intracellular 2-HG Quantities in Cell Lines

Intracellular 2-HG was measured following the kit manufacturer’s instructions. After cell dissociation, cells were lysed, then centrifuged. Deproteinization was performed by adding 4 M perchloric acid (PCA) to the supernatant. To neutralize the solution, KOH (2M) was added until the pH of the supernatant ranged from 6.5 to 8. For 2-HG detection, Resaruzin solution was added, and the plate was incubated at 37 °C for 45 minutes, protected from light before fluorescence measurement. For analysis, Blank wells’ values were subtracted from standards and samples.

### Telomere-FISH

Telomere-fluorescence in situ hybridization (Telo-FISH) was performed on metaphase spreads. Cells were cultured flasks treated with Karyomax for 2 hours, harvested using trypsin, and resuspended in hypotonic solution (0.6% sodium citrate) for 20 minutes at 37 °C. Cells were then fixed with ethanol/acetic acid (3V:1V) and stored overnight at 4°C. Metaphase chromosome preparations were obtained by spreading cells on SuperFrost microscope slides using a cytogenetic drying system (Thermotron AdGenix). After drying, Telo-FISH staining was performed using a Cy3-telomeric PNA probe (CCCTAA)_3_ (Applied Biosystem) as previously described[65]. Chromosomes were counterstained with DAPI. We used a Zeiss Axioplan2 microscope with Metafer imaging software to image metaphases. For each cell line, 80 to 208 metaphases were analyzed.

### Telomeric Repeat Amplification Protocol (TRAP)

Telomerase enzymatic activity was determined using the TRAPeze Telomerase detection kit according to the manufacturer’s instructions. Protein concentrations were determined using the Bradford method. Telomerase activity was assessed using 0.1 and 0.2 µg of proteins and TRAP products were separated on a 12.5% non-denaturing polyacrylamide gel electrophoresis (PAGE) in 0.5X TBE buffer. Gels were stained with 1 µg/ml of ethidium bromide and PCR amplification products were visualized under UV light exposure. To ensure reliability, an internal control band must be present in all lines on the PAGE, as samples may contain Taq polymerase inhibitors.

### Colocalization of PML/Telomeres and Detection of ALT-Associated PML Bodies (APBs)

Cells were cultured on Millicell^®^ EZ slides coated with human recombinant laminin, fixed with 4% PFA, and permeabilized with PBS, 0.5% Triton X-100. Blocking was performed in PBS, 7.5% fetal calf serum, 7.5% goat serum. Blocked cells were incubated overnight at 4° C with an α-PML antibody, and 1h at RT with an Alexa Fluor 488 α-rabbit secondary antibody. After incubation; slides were fixed in 4% formaldehyde, dehydrated in a graded ethanol solution (50%, 80%, and 100%) and incubated with a Cy3-telomeric PNA probe (CCCTAA)_3_ diluted in a hybridization solution (70% formamide, 10 mM Tris-HCl pH7.2, 1% BSA) at 80°C for 3 minutes, followed by hybridization for 2 hours at room temperature. After hybridization, slides were washed with buffer 1 (70% formamide, 10 mM Tris-HCl, pH 7.2), buffer 2 (50 mM Tris-HCl pH 7.2, 150 mM NaCl, and 0.05% Tween20), and PBS. Cells were counterstained with DAPI. After ethanol dehydration and air drying, slides were mounted using Fluoromount and stored overnight at 4° C. Images were captured using a Nikon A1 confocal microscope (objective x 60) and 29 to 168 cells were analyzed per astrocytoma.

### C-circle Assay

Amplification and detection of C-circles were performed as described by Henson[66], with minor modifications. DNA extraction was performed with QIAamp DNA Blood Mini Kit according to the manufacturer’s instructions, and DNA concentrations were determined using NanoDrop^TM^. Equal amounts of genomic DNA (30ng) were used in the rolling circle amplification with 7.5U of phi 29 DNA polymerase for 6h at 30°C. Telomeric qPCR was used to determine CCA levels by measuring the increase in total telomeric DNA produced during the assay and normalized to the total DNA amount determined by qPCR for the 36B4 gene. Each reaction included a negative control without phi 29 polymerase. For each sample, triplicate telomeric qPCR reactions and triplicate single-copy gene qPCR reactions (ribosomal lateral subunit gene P0, RLP0/36B4) were performed. Two independent experiments were conducted.

### Immunofluorescence

Cells seeded at a density of 30 000 cells/cm^2^ on coverslips coated with Poly-D-Lysine and Laminin were fixed with 4% paraformaldehyde (PFA) for 15 minutes at room temperature. Permeabilization and blocking were performed with 0.1% Triton and 5% donkey serum. Primary antibodies listed in **Table.S15** were incubated overnight at 4° C. Secondary antibodies Alexa 488, 574, or 647 (Jackson) and Hœchst 33342 were incubated for 1 hour at room temperature. Coverslips were mounted using Fluoromount. Images were taken using the Zeiss Apotome Z2 microscope. Image analyses were performed using Image J software. A minimum of 500 cells per experiment was counted to determine the proportion of each cell subpopulation.

### Flow Cytometry

After 5 days of culture with or without growth factors, cells were incubated with accumax for 30 minutes to detach them from the support, then centrifuged for 5 minutes at 1300 rpm at 4° C in cold PBS supplemented with 0.5% BSA. For extracellular staining, cells were incubated for 10 minutes at 4° C with the coupled antibodies (**Table.S16**). For intracellular staining, cells were fixed with 4% PFA for 15 minutes at 4° C. Permeabilization and blocking were performed with 0.1% saponin and 0.5% BSA. Primary antibodies listed in **Table.S16** were incubated for 30 minutes at 4°C. Secondary antibodies Alexa 488 or 647, and Hœchst 33342 were incubated for 30 minutes at 4°C. Flow cytometry recording was performed using the Macs Quant (Miltenyi) and analyses were performed using FlowJo software.

### Cell Proliferation Analysis by Dye Dilution

LGG275 cells were incubated with a CellTrace^TM^ solution before being seeded at a density of 25 000 cells/cm^2^ in a medium with or without growth factors for 20 days. For one of the conditions, after 10 days of culture without factors, EGF and FGF_2_ growth factors were added for an additional 10 days. Cells were then dissociated using the usual method and CellTrace^TM^ intensity was measured by flow cytometry (MACS Quant Analyzer).

### Cell Cycle Analysis

Cells were seeded at a density of 30,000 cells/cm^2^, then maintained in culture for 20 days in the presence or absence of growth factors, or 10 days without growth factors, then 10 days adding growth factors. For cell cycle analysis, cells were dissociated according to the previous protocol. 70% ethanol was added to cell pellets and stored at -20°C until the experiment. Samples were centrifuged for 2 minutes at 4 200 rpm, then incubated for 30 minutes with RNAse, DNAse ree, followed by incubation with propidium iodide for 10 minutes before analysis. Flow cytometry recording was performed using the MACS Quant and analyzed using FlowJo software.

### Lentivirus Infections

LGG275 cell line cells were infected with a luciferase-GFP virus or a DLL3-GFP virus to induce its overexpression. For the P27 reporter line, cells were infected with a control mVenus-BFP2 virus or a P27/mVenus-BFP2 virus with a multiplicity of infection (MOI) of 7. After cell amplification, they were purified based on GFP or BFP2 using the FACS ARIA sorter (**Table.S16**).

### Cell Sorting

Cells were seeded at a high concentration (> 130 000 cells/cm^2^) in a medium without growth factors. After 5 days of culture, cells were dissociated by adding Trypsin-EDTA 0.05X and incubating them for 30 minutes at 37°C. Then, cells were incubated for 10 minutes at 4° C in a cold PBS solution with 0.5% BSA and coupled antibodies (CD44-APC and GLAST-PE), then centrifuged for 5 minutes at 1,300 rpm at 4°C. To discriminate live cells from dead cells, SytoxBlue^TM^ was added before cell sorting. For LGG275 P27/mVenus-BFP2 cells, dissociation was performed in Trypsin-EDTA, 0.25X for 4 minutes. Cell sorting was performed with the FACS ARIA sorter. Cell subpopulations were seeded at a density of 30 000 cells/cm^2^ in the presence or absence of growth factors and cultured for 1, 6, 18, or 30 days, then fixed for experiments.

### Treatments of Subpopulations with DLL4 and γ-Secretase Inhibitors

After cell sorting, cell subpopulations were seeded at a density of 30 000 cells/cm^2^ on coverslips. For DLL4 treatment, coverslips were pretreated with PDL and then incubated overnight at 4°C with 0.1% BSA or DLL4. 30 minutes before the experiment, mouse laminin was added to each well. For treatment with γ-secretase inhibitors, coverslips were pretreated with PDL and mouse laminin. Cells were cultured without growth factors and treated every 24 hours with DMSO, DAPT or LY411575. Cells were maintained in culture for 6 days and then fixed for immunostaining experiments.

### RNA Extraction and qPCR

Cells were scraped on ice in cold PBS, then RNA was extracted using the RNeasy mini or micro kit according to the manufacturer’s instructions. 200 ng of cDNA were synthesized with random primers and GoScript^TM^ reverse transcriptase. Quantitative PCR was performed using 2 ng of cDNA in triplicate. Primers are listed in **Table.S18**. The KAPA SYBR^®^ FAST kit was used and qPCR reaction was performed on Light Cycler 480, (Roche). Gene expression was calculated using the Log(2^−ΔΔCt^) method, with β-actin (ACTB) as a housekeeping gene for normalization.

### Statistical Analyses

All experiments were performed at least three times for confirmation (unless otherwise indicated). Data are represented as mean ± standard error of the mean (SEM). Statistical differences in experiments were analyzed with the tests indicated in the legends using GraphPad 10 Prism software (Version 10.4.2, Inc La Jolla, CA, USA). Tests are performed in a paired manner. Significant differences between experimental groups were analyzed either by a student test or one-way ANOVA, followed by multiple comparison tests (Dunnett, Turkey).

## Data availability

Bulk and single cell RNA-seq datasets have been deposited in the NCBI Gene Expression Omnibus (GEO) under the following accession numbers:

Single-cell RNA sequencing datasets can be accessed at GSE263796. RNA sequencing of the 4 astrocytoma cell lines can be accessed at GSE308063, RNA sequencing of the CD44^+^GLAST^+^ and CD44^-^GLAST^-^ subpopulations can be accessed at GSE305451, RNA sequencing of mitotic (P27^-^) and quiescent (P27^+^) cells can be accessed at GSE306414, RNA sequencing of the effect of DLL3 overexpression can be accessed at GSE308910. Whole exome sequencing datasets can be accessed in the SRA database at PRJNA1345048.

## Language Editing Assistance

Stylistic improvements and clarity enhancements were assisted by ChatGPT (OpenAI) artificial intelligence–based language model, and by the corrector Antidote 12 (^©^2025 Druide informatique inc. - v2.0.1.

