## Supplemental figures and material_methods for "Multi-omics characterization of IDH-mutant astrocytoma-derived cell lines reveals NOTCH-regulated plastic quiescent astrocyte-like state"

Figure.S1

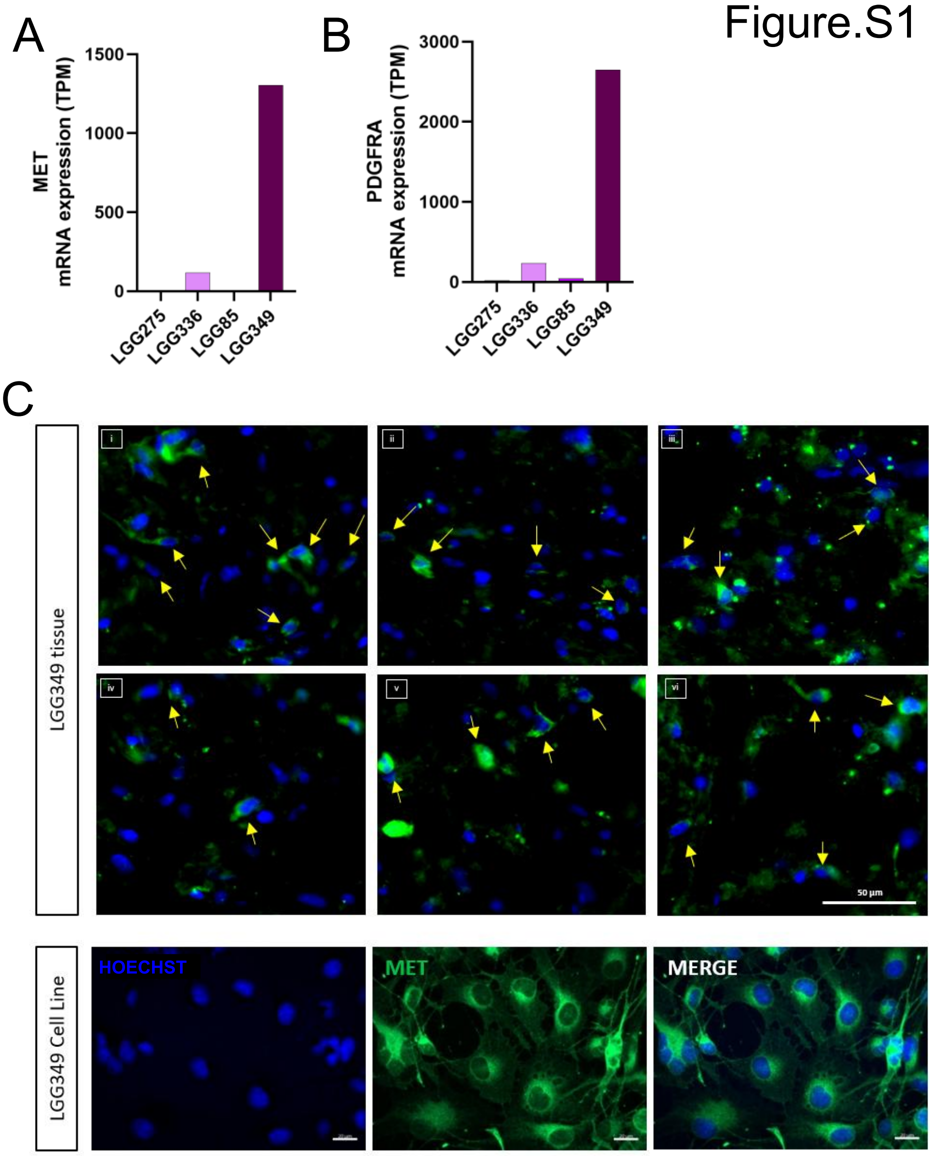
A
MET
mRNA expression (TPM)
1500
1000
500
0
LGG275
LGG336
LGG85
LGG349
B
PDGFRA
mRNA expression (TPM)
3000
2000
1000
0
LGG275
LGG336
LGG85
LGG349
C
LGG349 tissue
i
ii
iii
iv
v
vi
50 µm
LGG349 Cell Line
HOECHST
MET
MERGE

Figure.S2

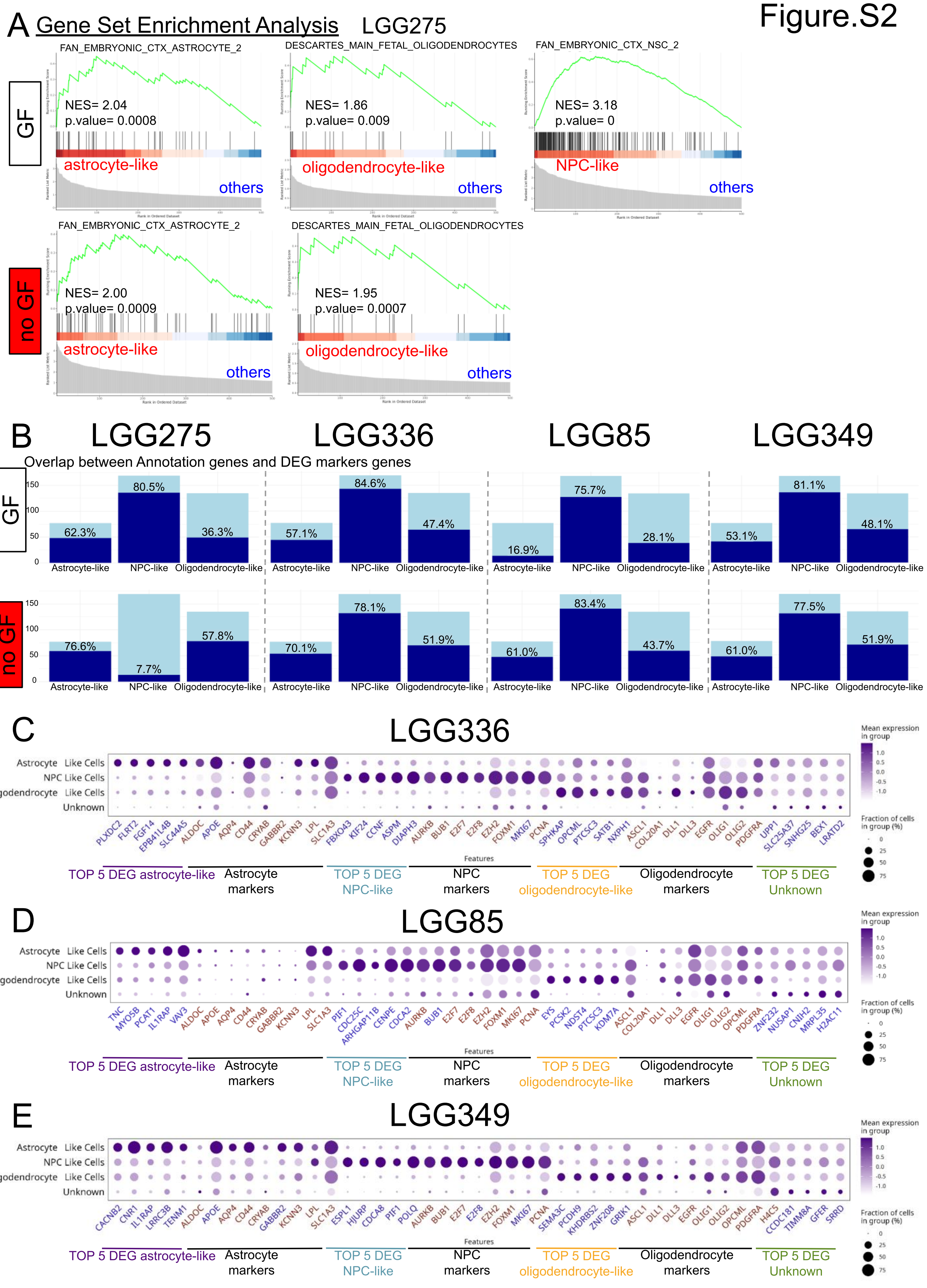

A Gene Set Enrichment Analysis LGG275
GF
no GF
FAN_EMBRYONIC_CTX_ASTROCYTE_2
NES= 2.04
p.value= 0.0008
astrocyte-like
others
DESCARTES_MAIN_FETAL_OLIGODENDROCYTES
NES= 1.86
p.value= 0.009
oligodendrocyte-like
others
FAN_EMBRYONIC_CTX_NSC_2
NES= 3.18
p.value= 0
NPC-like
others
FAN_EMBRYONIC_CTX_ASTROCYTE_2
NES= 2.00
p.value= 0.0009
astrocyte-like
others
DESCARTES_MAIN_FETAL_OLIGODENDROCYTES
NES= 1.95
p.value= 0.0007
oligodendrocyte-like
others
B LGG275 LGG336 LGG85 LGG349
Overlap between Annotation genes and DEG markers genes
GF
62.3% 80.5% 36.3%
57.1% 84.6% 47.4%
16.9% 75.7% 28.1%
53.1% 81.1% 48.1%
no GF
76.6% 7.7% 57.8%
70.1% 78.1% 51.9%
61.0% 83.4% 43.7%
61.0% 77.5% 51.9%
Astrocyte-like NPC-like Oligodendrocyte-like
C LGG336
D LGG85
E LGG349
Astrocyte Like Cells
NPC Like Cells
Oligodendrocyte Like Cells
Unknown
Identity
Features
Mean expression in group
Fraction of cells in group (%)
TOP 5 DEG astrocyte-like
Astrocyte markers
TOP 5 DEG NPC-like
NPC markers
TOP 5 DEG oligodendrocyte-like
Oligodendrocyte markers
TOP 5 DEG Unknown

Figure.S3

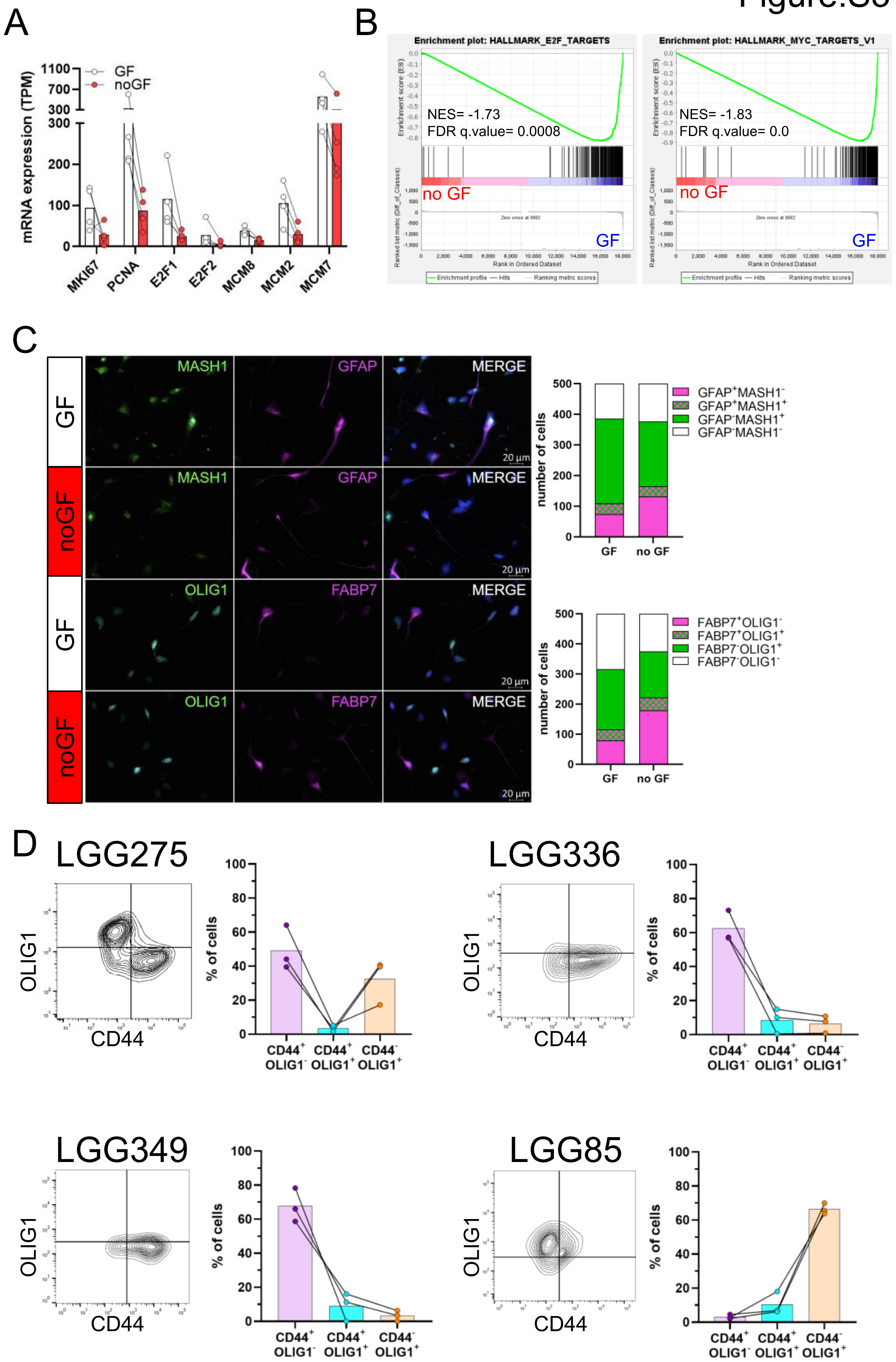
A
B
C
D
mRNA expression (TPM)
GF
noGF
MKI67
PCNA
E2F1
E2F2
MCM8
MCM2
MCM7
Enrichment plot: HALLMARK_E2F_TARGETS
Enrichment plot: HALLMARK_MYC_TARGETS_V1
NES= -1.73
FDR q.value= 0.0008
NES= -1.83
FDR q.value= 0.0
no GF
GF
MASH1
GFAP
MERGE
OLIG1
FABP7
20 µm
number of cells
GFAP+MASH1-
GFAP+MASH1+
GFAP-MASH1+
GFAP-MASH1-
FABP7+OLIG1-
FABP7+OLIG1+
FABP7-OLIG1+
FABP7-OLIG1-
LGG275
LGG336
LGG349
LGG85
OLIG1
CD44
% of cells
CD44+ OLIG1-
CD44+ OLIG1+
CD44- OLIG1+

Figure.S4

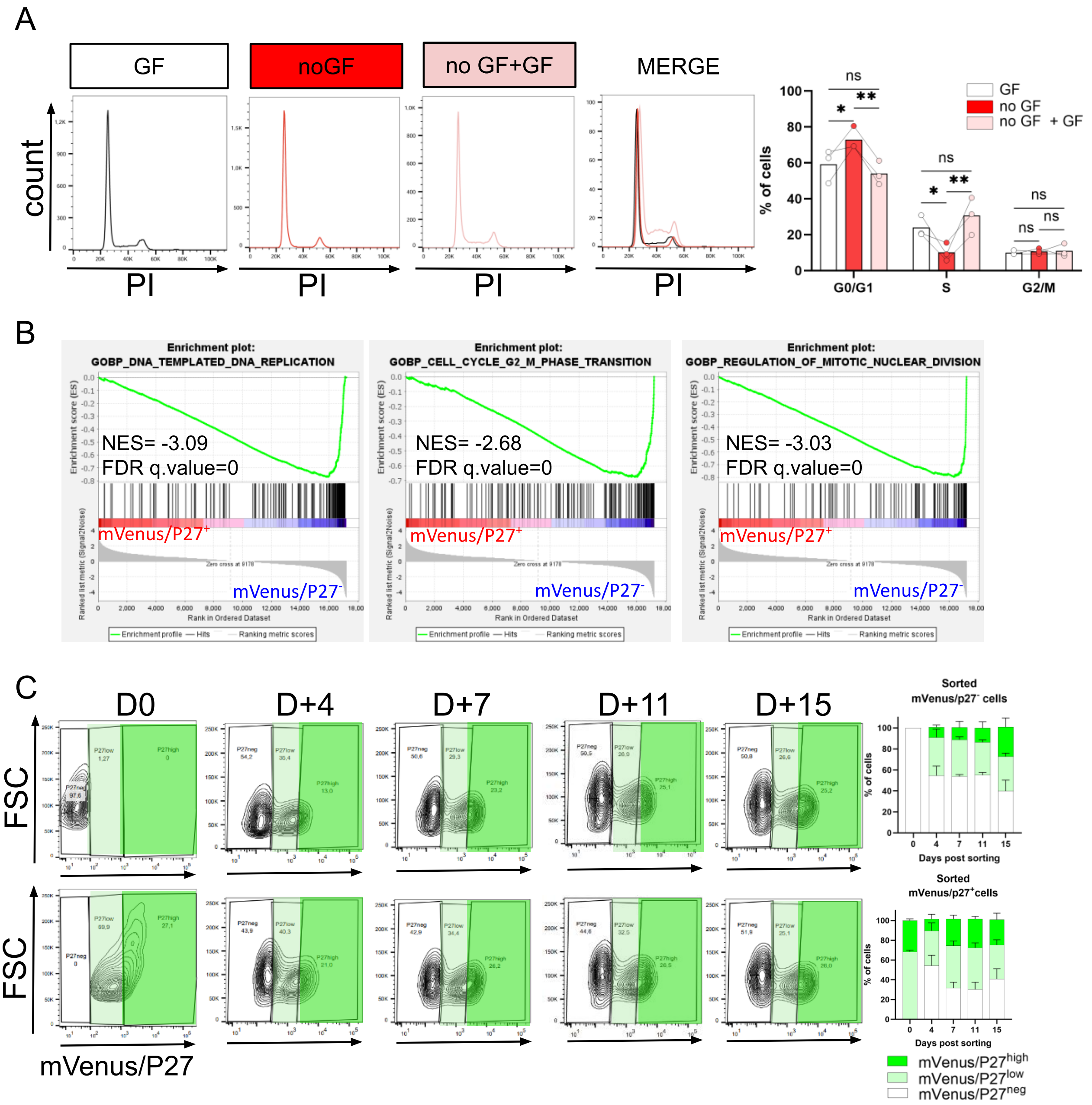
A
GF
noGF
no GF+GF
MERGE
count
PI
% of cells
GF
no GF
no GF + GF
ns
*
**
G0/G1
S
G2/M
B
Enrichment plot:
GOBP_DNA_TEMPLATED_DNA_REPLICATION
NES= -3.09
FDR q.value=0
Enrichment plot:
GOBP_CELL_CYCLE_G2_M_PHASE_TRANSITION
NES= -2.68
FDR q.value=0
Enrichment plot:
GOBP_REGULATION_OF_MITOTIC_NUCLEAR_DIVISION
NES= -3.03
FDR q.value=0
mVenus/P27+
mVenus/P27-
Enrichment score (ES)
Ranked list metric (Signal2Noise)
Rank in Ordered Dataset
Enrichment profile — Hits
Ranking metric scores
C
D0
D+4
D+7
D+11
D+15
mVenus/p27- cells
mVenus/p27+ cells
FSC
mVenus/P27
Sorted mVenus/p27- cells
Sorted mVenus/p27+cells
% of cells
Days post sorting
mVenus/P27high
mVenus/P27low
mVenus/P27neg

Figure.S5

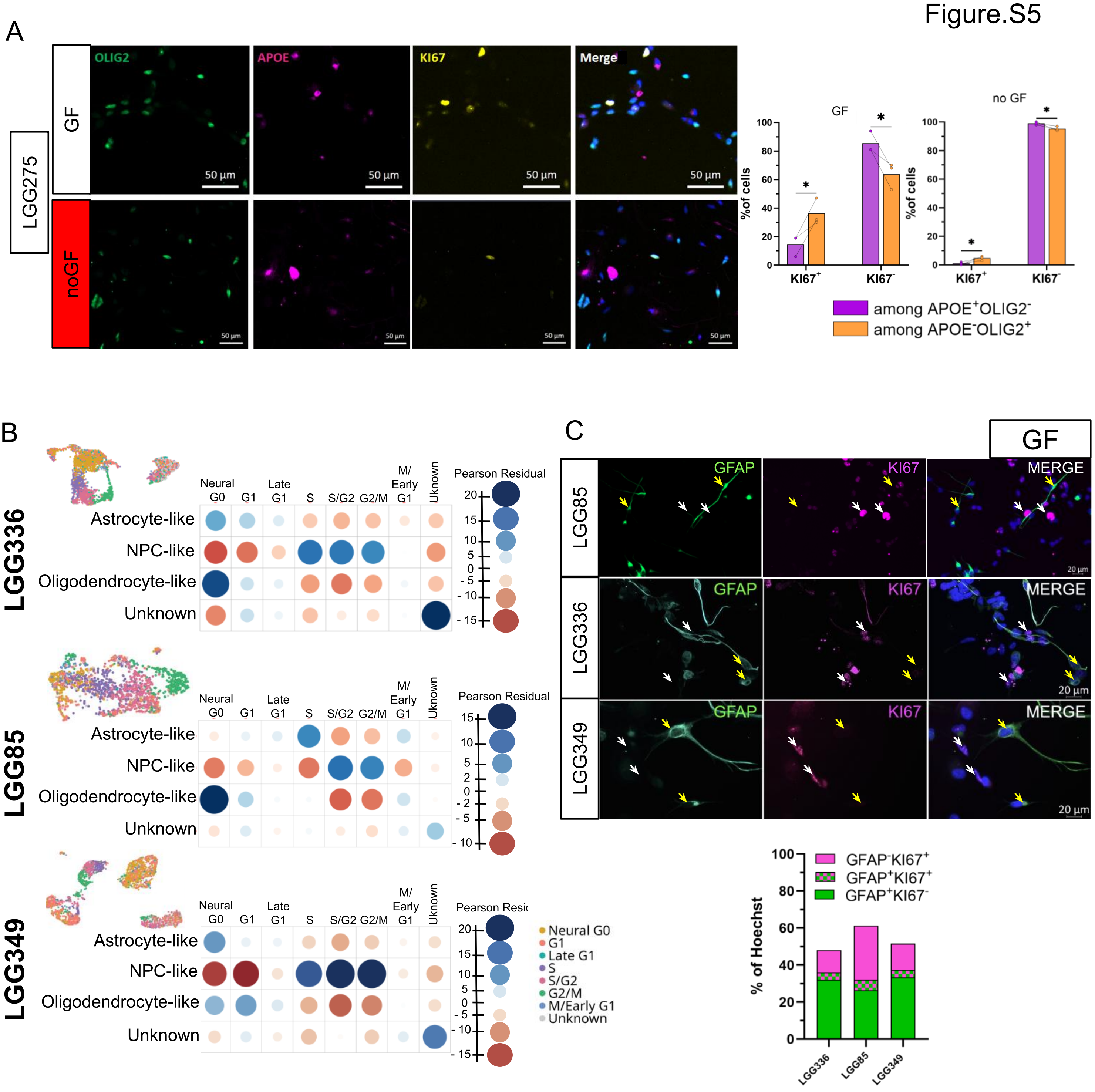
A
LGG275
GF
noGF
OLIG2
APOE
KI67
Merge
50 µm
%of cells
no GF
among APOE+OLIG2-
among APOE-OLIG2+
B
LGG336
LGG85
LGG349
Neural G0
G1
Late G1
S
S/G2
G2/M
M/ Early G1
Uknown
Astrocyte-like
NPC-like
Oligodendrocyte-like
Unknown
Pearson Residual
Neural G0
G1
Late G1
S
S/G2
G2/M
M/Early G1
Unknown
C
GF
GFAP
KI67
MERGE
20 µm
% of Hoechst
GFAP-KI67+
GFAP+KI67+
GFAP+KI67-

Figure.S6

A

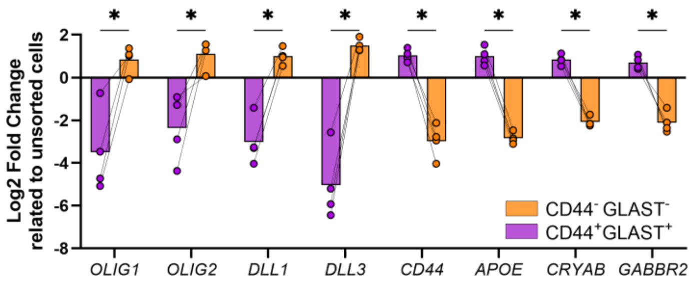


B

RNA

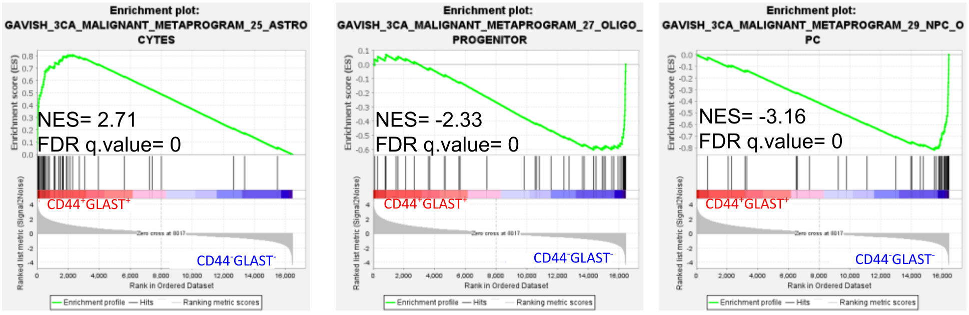

Figure.S7

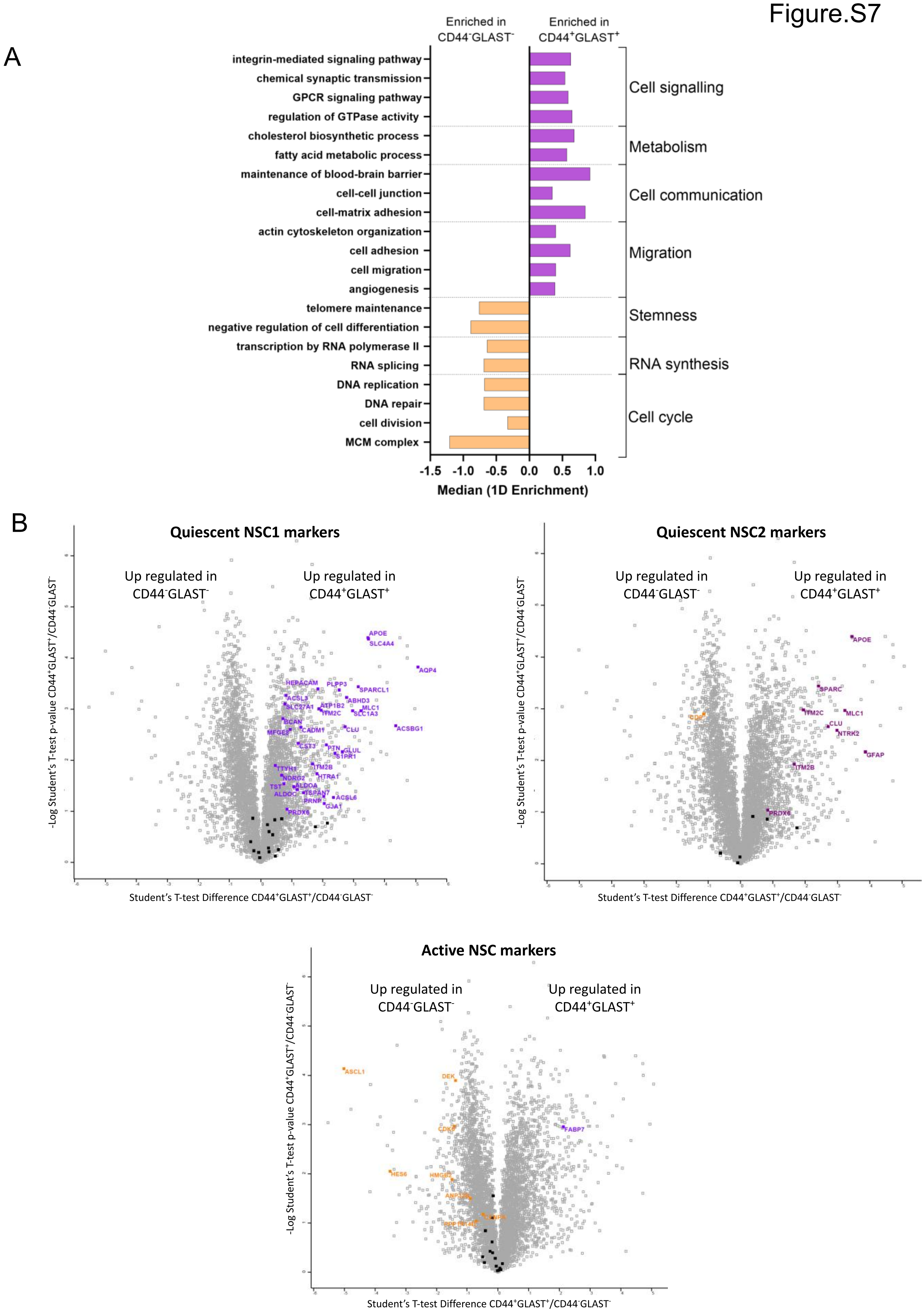
A
Enriched in CD44⁻GLAST⁻
Enriched in CD44⁺GLAST⁺
integrin-mediated signaling pathway
chemical synaptic transmission
GPCR signaling pathway
regulation of GTPase activity
cholesterol biosynthetic process
fatty acid metabolic process
maintenance of blood-brain barrier
cell-cell junction
cell-matrix adhesion
actin cytoskeleton organization
cell adhesion
cell migration
angiogenesis
telomere maintenance
negative regulation of cell differentiation
transcription by RNA polymerase II
RNA splicing
DNA replication
DNA repair
cell division
MCM complex
Cell signalling
Metabolism
Cell communication
Migration
Stemness
RNA synthesis
Cell cycle
-1.5 -1.0 -0.5 0.0 0.5 1.0
Median (1D Enrichment)
B
Quiescent NSC1 markers
Up regulated in CD44⁻GLAST⁻
Up regulated in CD44⁺GLAST⁺
APOE
SLC4A4
AQP4
HEPACAM
PLPP3
SPARCL1
ABHD3
MLC1
SLC1A3
ACSBG1
CLU
GLUL
PTN
TTYH1
ITM2B
HTRA1
NDRG2
TST
ALDOC
TSPAN7
PRNP
ACSL6
GJA1
PRDX6
-Log Student's T-test p-value CD44⁺GLAST⁺/CD44⁻GLAST⁻
Student's T-test Difference CD44⁺GLAST⁺/CD44⁻GLAST⁻
Quiescent NSC2 markers
Up regulated in CD44⁻GLAST⁻
Up regulated in CD44⁺GLAST⁺
APOE
SPARC
ITM2C
MLC1
CLU
NTRK2
GFAP
ITM2B
PRDX6
-Log Student's T-test p-value CD44⁺GLAST⁺/CD44⁻GLAST⁻
Student's T-test Difference CD44⁺GLAST⁺/CD44⁻GLAST⁻
Active NSC markers
Up regulated in CD44⁻GLAST⁻
Up regulated in CD44⁺GLAST⁺
ASCL1
DEK
FABP7
HES6
-Log Student's T-test p-value CD44⁺GLAST⁺/CD44⁻GLAST⁻
Student's T-test Difference CD44⁺GLAST⁺/CD44⁻GLAST⁻

Figure.S8

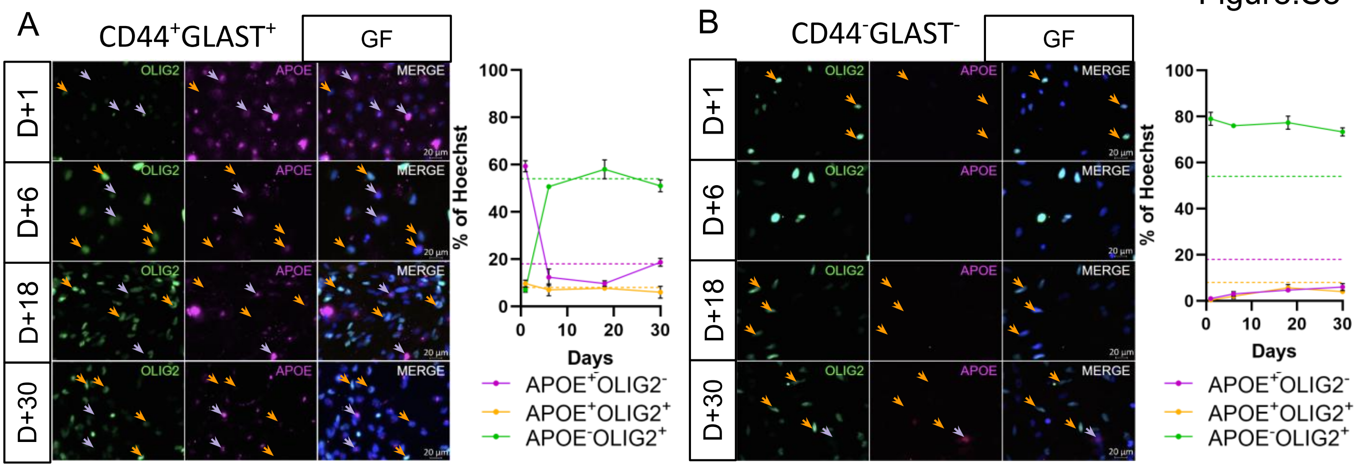

A
CD44+GLAST+
GF
OLIG2
APOE
MERGE
D+1
D+6
D+18
D+30
20 µm
% of Hoechst
Days
APOE+OLIG2-
APOE+OLIG2+
APOE-OLIG2+
B
CD44-GLAST-

Figure.S9

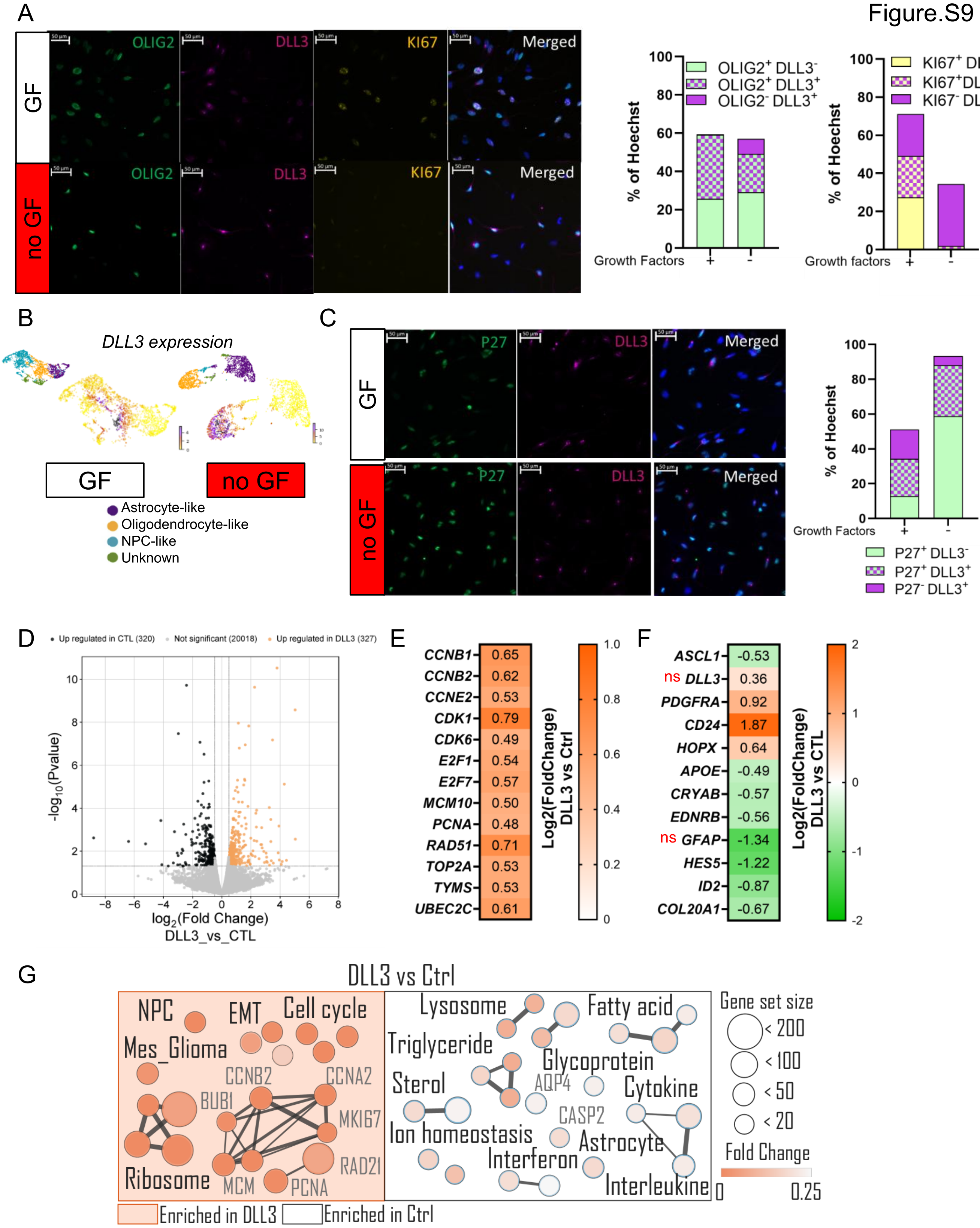
A
GF
no GF
OLIG2
DLL3
KI67
Merged
50 µm
% of Hoechst
Growth Factors
+
-
OLIG2+ DLL3-
OLIG2+ DLL3+
OLIG2- DLL3+
Growth factors
KI67+ DLL3-
KI67+DLL3+
KI67- DLL3+
B
DLL3 expression
GF
no GF
Astrocyte-like
Oligodendrocyte-like
NPC-like
Unknown
C
GF
no GF
P27
DLL3
Merged
% of Hoechst
Growth Factors
P27+ DLL3-
P27+ DLL3+
P27- DLL3+
D
Up regulated in CTL (320)
Not significant (20018)
Up regulated in DLL3 (327)
-log10(Pvalue)
log2(Fold Change)
DLL3_vs_CTL
E
CCNB1 0.65
CCNB2 0.62
CCNE2 0.53
CDK1 0.79
CDK6 0.49
E2F1 0.54
E2F7 0.57
MCM10 0.50
PCNA 0.48
RAD51 0.71
TOP2A 0.53
TYMS 0.53
UBEC2C 0.61
Log2(FoldChange) DLL3 vs Ctrl
F
ASCL1 -0.53
ns DLL3 0.36
PDGFRA 0.92
CD24 1.87
HOPX 0.64
APOE -0.49
CRYAB -0.57
EDNRB -0.56
ns GFAP -1.34
HES5 -1.22
ID2 -0.87
COL20A1 -0.67
Log2(FoldChange) DLL3 vs CTL
G
DLL3 vs Ctrl
NPC
EMT
Cell cycle
Mes_Glioma
CCNB2
CCNA2
BUB1
MKI67
RAD21
Ribosome
MCM
PCNA
Lysosome
Fatty acid
Triglyceride
Glycoprotein
Sterol
AQP4
Cytokine
CASP2
Ion homeostasis
Astrocyte
Interferon
Interleukine
Gene set size
< 200
< 100
< 50
< 20
Fold Change
0
0.25
Enriched in DLL3
Enriched in Ctrl

Figure.S10

A

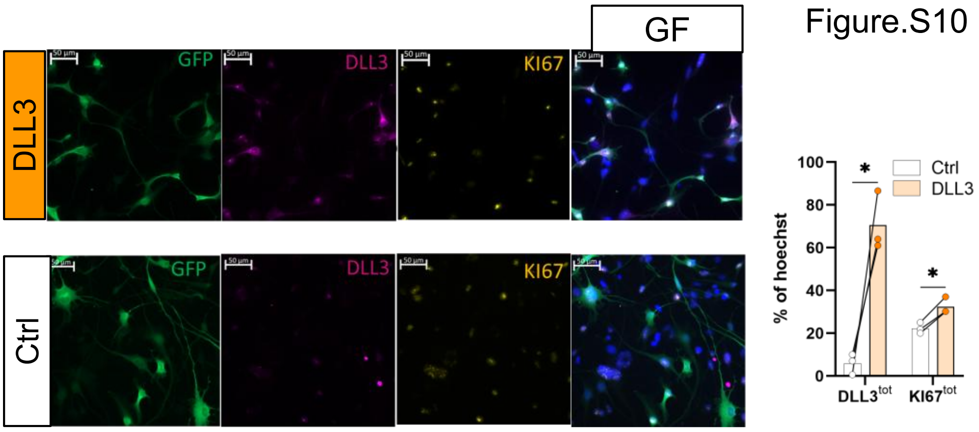


B

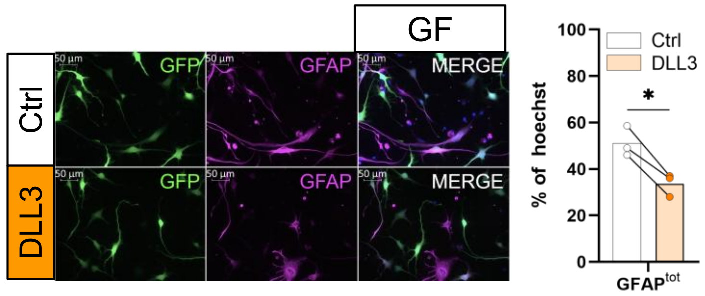

**Figure S1: Altereted gene expression in the 4 astrocytoma cell lines**
(A) *MET* and (B) de *PDGFRA* mRNA expression (TPM = Transcript Per Million) in LGG275, 336, 85 and 349 (n=1) (C) IF analysis of MET expression in 349 tumor resection and LGG349 cell line, scale bar = 50µm for tissue, 20µm for cell line.

**Figure S2: scRNA sequencing and RNA sequencing analysis of 4 astrocytoma cell lines**
(A) GSEA analysis showing enrichment of astrocyte, oligodendrocyte and NPC signatures in astrocyte-like, oligodendrocyte-like and NPC-like subpopulation, respectively, in LGG275 cell lines, with or without (growth factors. P.value was calculated using Benjamini-Hochberg methods. NES = Normalized Enrichment Score, FDR = False Discovery Rate (B) Histogram showing significant DEG (dark) blue among reference (light) astrocyte, oligodendrocyte and NPC gene list (C) Dot Plot of astrocyte, oligodendrocyte and proliferative markers expression in each subpopulation of LGG336 (E) LGG85 and (F) LGG349 with growth factors.

**Figure S3: Astrocyte and oligodendrocyte markers represent two non-overlapping subpopulations in astrocytomas cell lines** (A) mRNA expression of proliferative genes with and without GF in the 4 astrocytomas cell lines. Bars represent the mean expression and each dots represents one cell line (LGG275, LGG336, LGG349 et LGG85, n=4) (B) GSEA analysis of RNA sequencing comparing 4 atsrocytoma cell lines cultured with (GF) versus without GF (noGF) shows downregulation of proliferative signatures after GF withdraw (C) Analysis by IF of astrocyte (GFAP, FABP7) and oligodendrocyte (OLIG1, MASH1) markers, showing astrocyte-like and oligodendrocyte/NPC-like subpopulation in LGG275 cell line with (GF) or without GF (noGF), scale bar = 20µm. Left panel: Barplot show quantification as number of cells, (n=1, count = 500 cells) (D) CD44 and OLIG1 expression by flow cytometry showing astrocyte-like (purple) and oligodendrocyte/NPC-like (orange) population in the 4 astrocytoma cell lines without growth factor. Barplot show quantification as % of cells (Bars = mean, dots = n = 3).

**Figure S4: There is a subpopulation of quiescent cells within LGG275 cell line**
(A) Cell cycle analysis by flow cytometry using propidium iodide in LGG275 cell line after 20 days with (GF) or without (no GF) GF or 10 days without GF following by 10 days with GF (no GF + GF). Right Panel: Barplot show Barplot show quantification as % of cells (bars = mean, dots = n = 3) (ns = non-significant, * p<0, 05, **<0,01, multiple comparison Turkey test) (B) GSEA analysis of RNA sequencing comparing mVenus/P27$^+$ cells versus mVenus/P27$^-$ cells shows a enrichment of cell cycle gene signatures in mVenus/P27$^-$ cells in LGG275 line. Q.value has been calculated using Benjamini-Hochberg method. NES = Normalized Enrichment Score, FDR = False Discovery Rate. (C) Flow cytometry analysis of mVenus/P27 fusion protein expression 0, 4, 7, 11 et 15 days in mVenus/P27$^+$ and mVenus/P27$^-$ populations. Right panel: Barplot show quantification of negative, low and high mVenus/P27 population as % of cells (bars = mean ± S.E.M, n=3).

**Figure S5: LGG275, LGG336 and LGG349 cell lines have quiescent cells with a major astrocyte-like phenotype, followed by an oligodendrocyte-like phenotype**
(A) IF analysis of APOE, OLIG2 and KI67 expression in LGG275 line, identifying KI67 preferential expression in oligodendrocyte/NPC-like (APOE$^-$OLIG2$^+$) rather than in astrocyte-like cells (APOE$^+$OLIG2$^-$) with and without GF, scale bar = 50µm. Right panels: Barplot show quantification as % of Hoechst cells (bars = mean, dots = n = 3) (* p<0, 05, multiple comparison Turkey test) (B) UMAP colored by ccAFv2 cell-cycle state classifier, including Neural G0. Right, dot plot showing the association between phenotypic states (astrocyte-like, NPC-like, oligodendrocyte-like, unknown) and ccAFv2 cell-cycle classes (Pearson residual) in LGG336,

LGG85 and LGG349 lines. Significance was assessed by a $\chi^2$ test ($p < 0.01$) (C) IF analysis of GFAP and KI67 expression grown with GF showing two non-overlapping astrocyte-like ($GFAP^+$, yellow arrows) and mitotic ($KI67^+$, white arrow) populations in LGG85, LGG336 and LGG349 cell lines, scale bar = 20µm. Below: Barplot show quantification as % of $Hoechst^+$ cells (n=1, count = 500 cells).

**Figure S6 : Cell sorting allows efficient isolation of "astrocyte-like" and "oligodendrocyte-like" cells**
(A) Log2(FoldChange) of astrocyte and oligodendrocyte markers expression in $CD44^+GLAST^+$ (purple) and $CD44^-GLAST^-$ (orange) populations. Unsorted cells were used as control to calculate the Log2(FoldChange) (bars = mean, dots = n = 3) (* p<0,05, multiple Student test) (B) GSEA analysis of RNA sequencing comparing $CD44^+GLAST^+$ versus $CD44^-GLAST^-$ populations shows enrichment of astrocyte signatures in $CD44^+GLAST^+$ cells, and OPC and NPC metaprogram signatures in $CD44^-GLAST^-$ cells (n=3). NES = Normalized Enrichment Score, FDR = False Discovery Rate. P.value has been calculated using the Benjamini-Hochberg method.

**Figure S7: Proteomic analysis confirms transcriptomic results of the two populations** (A) 1D enrichment and fisher analyses in $CD44^+GLAST^+$ versus $CD44^-GLAST^-$ cells (n=3). Bars represent the median enrichment. All signaling pathways are significant a p.value <0,01 as determined by a multiple Student t-test (B) Volcano Plot of proteins differentially expressed associated with quiescent NSC1, NSC2 and active NSC phenotype in $CD44^+GLAST^+$ versus $CD44^-GLAST^-$ cells (filled square). Orange and purple dots are significant differentially expressed proteins (Student t-test, permutation based FDR, FDR = 0.05, s0 = 0.1).

**Figure S8: Astrocyte-like cells are plastics and can transit into an "oligodendrocyte/NPC-like" state with growth factors** (A) IF analysis of APOE and OLIG2 expression in the LGG275 cell line, identifying astrocyte-like ($APOE^+OLIG2^–$) cells (purple arrows) and oligodendrocyte/NPC-like ($APOE^–OLIG2^+$) cells (orange arrows) with GF, in $CD44^+GLAST^+$ and (B) $CD44^-GLAST^-$ populations, 1, 6, 18, and 30 days after cell sorting, scale bar = 20µm. Barplot show quantification as % of $Hoechst^+$ cells (dots on curve = mean ± S.E.M, n=3)

**Figure S9: Overexpression of DLL3 leads to a decrease in the astrocyte-like phenotype in favor of oligodendrocyte-like and NPC-like phenotypes in the LGG275 cell line**
(A) IF analysis of OLIG2, DLL3 and KI67 expression in LGG275 line with or without GF, showing expression pattern of DLL3 in oligodendrocyte/NPC-like ($OLIG2^+$) and $OLIG2^-$ populations, as well as KI67 expression in $DLL3^+$ cells, scale bar = 50µm. Right panel; Barplot show quantification as % of $Hoecsht^+$ cells (n=1, count = 500 cells) (B) RNA expression of *DLL3* in LGG275 using scVelo on scRNA sequencing data (C) IF analysis of P27 and DLL3 expression in LGG275 line with or without GF, showing $DLL3^+$ quiescent ($P27^+$) or non-quiescent ($P27^-$) cells, scale bar = 50µm. Right panel: Barplot show quantification as % of $hoecsht^+$ cells (n=1, count = 500 cells) (D) VolcanoPlot of DEG between DLL3 overexpression vs Ctrl (E) Log2(FoldChange) of proliferative markers between DLL3 overexpression vs Ctrl (F) Log2(FoldChange) of oligodendrocyte, stemness and astrocyte markers between DLL3 overexpression vs Ctrl. All genes are differentially expressed with p<0.05, except those with ns (non-significant) mention. (G) Cytoscape visualization of enrichment signaling pathways between DLL3 overexpression vs Ctrl. P.value<0.05 and FDR q.value < 0.25 as determinated by the Benjamini-Hochberg method (n=3).

**Figure S10: DLL3 overexpression leads a decrease of "astrocyte-like" GFAP$^+$ in LGG336 cell lines**

(A) IF analsyis of DLL3 and KI67 expression in LGG336 cell line, with GF, showing DLL3$^+$ (white arrows) and mitotic KI67$^+$ (yellow arrow) states proportions after DLL3 overexpression Right panel: Barplot show quantification as % of Hoeschst$^+$ cells (bars = mean, dots = n = 3) (*p<0.05, Dunnett's multiple comparison test). (B) IF analysis of GFAP expression in LGG336 line with GF, following astrocyte-like state proportion after DLL3 overexpression. Right panel: Barplot show quantification as % of Hoeschst$^+$ cells (bars = mean, dots = n = 3) (*p<0.05, Dunnett's multiple comparison

### Supplemental Material and methods

#### Whole Exome Sequencing

DNA extraction from the four astrocytoma cell lines was performed using Wizard Genomic DNA Purification kit. DNA extracts were then sent to the DNBSEQ BGI platform for sequencing. BGI supervised all procedures, including sample preparation, sequencing, and bioinformatics analyses. Raw data were cleaned using SOAPnuke filter. Clean datas were then aligned using the human reference genome GRCh38-p13, Burrows-Wheeler Aligner (BWA V0.7.17 - http://bio-bwa.sourceforge.net/) and BWA-MEM method. Duplicate reads were removed using GATK MarkDuplicates tool (v4.1.4.1, https://gatk.broadinstitute.org/hc/en-us/articles/360037225972-MarkDuplicates). Base quality values have been improved using GATK BaseRecalibrator (v4.1.4.1, https://gatk.broadinstitute.org/hc/en-us/articles/360037593511-BaseRecalibrator) and GATK ApplyBQSR (v4.1.4.1, http://bio-bwa.sourceforge.net/) softwares. To detect known mutations and SNP in each cell line, we used VarDecrypt software, developed by Salma and al(1).

#### RNA Sequencing

For each experiment, RNA samples were extracted, as described previously. RNA was sent to the BGI Genomics DNBSEQ platform (Poland or Hong Kong - China). RNA library construction, sequencing, quality control, and bioinformatics analysis were all performed by the platform. Library preparation was performed using an Optimal Dual-mode mRNA Library Prep Kit. RNA was denatured at a suitable temperature to open the secondary structure, and mRNA is enriched by oligo (dT) attached magnetic beads. After reacting at a suitable temperature for a fixed time period, RNAs were fragmented with fragmentation reagents. Then first-strand cDNA was generated using random hexamer-primed reverse transcription with the fragmented RNA as a template. To synthesize the second-strand cDNA, the synthesis reaction system was prepared and dUTP is used to replace dTTP. After obtaining the double strand cDNA product, it was converted to a blunt end with an end repair reaction. After cDNA end repair, a single ‘A’ nucleotide was added to the 3’ ends of the blunt fragments through A tailing reaction. Then the library adapters were ligated to the two ends of the cDNA with a ligation reaction. Finally, the library products were amplified through PCR reaction and subjected to quality control. Next, the single-stranded library products were produced via denaturation. The reaction system for circularization was set up to get the single-stranded circularized DNA products. Any single stranded linear DNA molecules were digested. The final single-stranded circularized library was amplified by phi29 and rolling circle amplification (RCA) to make a DNA nanoball (DNB), which carries more than 300 copies of the initial single-stranded circularized library molecule. The DNBs were loaded into the patterned nanoarray and sequencing reads with PE 150 bases length are generated on the DNBSEQ-T7 platform, generating 20M paired reads per sample. Raw sequencing data were filtered using SOAPnuke (v1.5.2), a quality control software independently developed by BGI. The following parameters were used: -l 15, -q 0.2, and -n 0.05. RNA-seq data were analyzed using the DESeq2 package. Volcano plots and bubble charts were generated using the SRPlot web tool (https://www.bioinformatics.com.cn).

**Gene Set Enrichment Analysis (GSEA)**

The GSEA analysis software (Version 4.3.3) was used to perform a comparative analysis of differentially expressed genes and molecular profiles between different conditions after RNA sequencing. The compared gene lists are the "normalized read count". The gene ranking method is the "Signal2Noise" method for groups consisting of triplicates and "Diff of class" for groups com+ed of duplicates. The collections used for comparison in this paper are the "Curated c2" collections (c2.all. v2024.1.Hs.symbols.gmt), "Cell types c8" (c8.all. v2024.1.Hs.symbols.gmt), "Gene Ontology c5" (c5.all. v2024.1.Hs.symbols.gmt) and "Computational c4" (c4.all. b2024.Hs.symbols.gmt). The "quiescent NSC1" (qNSC1), qNSC2 and "active NSC" (aNSC) gene lists were established from Kalamakis et al(2). The p.value was calculated using the Benjamini-Hochberg method. NOTCH_UP and NOTCH_DOWN genelist were constructed by treated cells with DLL4, an activator of NOTCH, then isolate significant up (NOTCH_UP) and down (NOTCH_DOWN) regulated genes following NOTCH activation (Table.S18).

**Cytoscape Mapping**

Visualization of GSEA analyses related to the gene ontology collection c5 is performed using Cytoscape software (Version 3.3) by filtering the results and keeping only data with a p.value less than 0.05 and FDR q.value less than 0.25, with an overlap coefficient less than 0.5. This visualization highlights genes common to different signaling pathways.

**Single-Cell 3' RNA Sequencing Using the Chromium Platform**

*Cell Preparation*

Cells were seeded at a density of 200,000 cells/cm$^2$ in a flask pretreated with polyHEMA for the condition with growth factors. For the condition without growth factors, cells were seeded at a density of 40,000 cells/cm$^2$ in a flask pretreated with PDL and murine laminin. After 4 days of culture with or without growth factors, neurospheres were collected by centrifugation for 1 minute at 1300 rpm at room temperature. Then, cells were dissociated following the usual protocol. To remove aggregates, the cell suspension was filtered through a 40 µm sieve using cold PBS, then centrifuged for 5 minutes at 1,300 rpm at 4° C. To remove debris, cell pellets were resuspended in a 15% Percoll solution and centrifuged for 5 minutes at 1,300 rpm at 4° C, then rinsed with a PBS solution containing 0.04% BSA. The final cell suspension was prepared in a PBS solution containing 0.04% BSA with a final concentration of 1000 cells/µL.

*Library Preparation and Sequencing*

Cell suspensions were loaded onto a Chromium controller (10x Genomics, Pleasanton, CA) to generate single-cell Gel Beads-in-Emulsion (GEMs). Single-cell RNA-Sequencing libraries were prepared using the Chromium Next GEM Single Cell 3' V3.1 reagent kits (Dual Index, P/N 1000268, 10x Genomics). Briefly, reverse transcription was performed at 53°C for 45 minutes, followed by incubation at 85°C for 5 minutes. GEMs were then broken and single-strand cDNAs were cleaned with DynaBeads MyOne Silane (Thermo Fisher Scientific; P/ N 37002D). cDNAs were amplified by PCR, cleaned with SPRIselect beads (SPRI P/N B23318), fragmented, end-repaired, A-tailed, and size-selected with SPRIselect beads. Indexed adapters were ligated and cleaned with SPRIselect beads. The resulting DNA fragments were amplified by PCR and size-selected with SPRIselect beads. The size distribution of the resulting libraries was monitored using a fragment analyzer (Agilent Technologies, Santa Clara, CA, USA) and libraries were quantified using the KAPA library quantification kit (Roche, Basel, Switzerland). Libraries were denatured with NaOH, neutralized with Tris-HCl, and diluted to 150 pM. Clustering and sequencing were performed on a NovaSeq 6000 (Illumina, San Diego, CA, USA) using the paired-end 28-90 nt protocol on an SP flow cell lane and an S4 flow cell lane 8.

*Alignment, Processing, and Quality Control of ScRNAseq Data*

Preprocessing of single-cell RNA sequencing (scRNA-sequencing) data was performed using the 10x Genomics Cell Ranger v7.1.0 pipeline. Raw sequencing data were processed with “cellranger mkfastq” to generate FASTQ files, followed by “cellranger count” for alignment, filtering, and UMI counting against the human genome GRCh38-p13. The resulting count matrices were imported into R using the Read10X function from the Seurat R package (v5.2.0). Eight LGG scRNAseq samples were included in our analysis. Seurat objects were created for each sample, with cells containing fewer than 200 genes or more than 20% mitochondrial reads filtered out. Gene expression was normalized using the LogNormalize method, and the 2000 most variable genes were identified using the FindVariableFeatures function. Principal component analysis (PCA) was performed on these variable genes, followed by UMAP dimensionality reduction using the top 30 principal components. Clustering was performed using the FindNeighbors and FindClusters functions.

*Annotation*

Cell type annotation was performed using a multi-step approach combining the expression of known marker genes, transcription factor expression analysis, co-expression motifs, and ScType (a computational method for fully automated and ultra-fast cell type identification based solely on the provided scRNA-seq data)(3). We first analyzed the expression of lineage-specific markers for oligodendrocytes and OPCs, astrocytes, and NPCs using Seurat's FeaturePlots and ViolinPlots. We also collected human gene lists for OPCs and oligodendrocytes and astrocytes from CellMarker 2.0 (Copyright© College of Bioinformatics Science and Technology, Harbin Medical University)(4). The lists were then sorted to keep only cell type-specific genes using The Human Protein Atlas database (Version 23.0 Version 23.0 –(https://www.proteinatlas.org/). The NPC-like list was constructed from the list established by Zhong et al(5) and supplemented by the list from Tirosh et al(6). The AddModulescore function from Seurat was used to calculate a gene list score, which we then visualized using Seurat's FeaturePlots and ViolinPlots. From these steps, we built a comprehensive set of cell type markers. This complete gene list was used with the ScType algorithm to score and classify cells into 4 categories: astrocyte-like, oligodendrocyte-like, NPC-like, and unknown (cells with a ScType score <0). The proportion of annotated cells for each putative cell type was quantified and visualized using pie charts. To validate the annotations, we performed differential gene expression analysis between classified cell populations and conducted GSEA analysis using the C8 collection from the MSigDB database. Enrichment of cell type-specific pathways in differentially expressed genes supported the accuracy of our annotations.

*Cell Cycle*

Cell cycle analysis was performed using the ccAFv2 algorithm (Cell cycle classifier for R and Seurat)(7) to classify cells into six cell cycle states (G1, G1 late, S, S/G2, G2/M, and M/G1 early) and a Neural G0-like state. The default parameter of this classifier was applied to the Seurat object of all samples. To evaluate the correlation between cell phenotype and cell cycle phase as well as frequency, Pearson residuals were calculated and the p.value were determined by a Chi2 test.

*Intercellular Communication*

To elucidate intercellular communication patterns within the samples, we used CellChat v2.1.2(8), a computational tool to infer and analyze intercellular communication networks from single-cell RNA sequencing data. We prepared the required input data from our Seurat object and used the CellChatDB. Human database for ligand-receptor interactions. The probability of intercellular communication was inferred using a mass action model, integrating gene expression data with prior knowledge of signaling interactions. We applied the default “trimean” method for robust estimation of mean gene expression, prioritizing stronger interactions for subsequent experimental validation. Communication networks were calculated at both individual ligand-receptor pair and signaling pathway levels. We visualized the aggregated intercellular communication network using chord diagrams to represent interaction counts and strengths between cell groups. Heatmaps were generated to identify signals contributing most to outgoing or incoming signaling of specific cell types. To discover coordinated communication patterns, we performed matrix factorization on outgoing and incoming intercellular communication probabilities, visualizing the results with river plots and dot plots. Finally, we used chord diagrams to visualize intercellular communication mediated by multiple ligand-receptor pairs or signaling pathways, both globally and for each cell type.

**RNA Velocity and Cellular Trajectory Analysis**

To infer cellular dynamics and differentiation trajectories, we used RNA velocity analysis with the velocity and scVelo packages. Initially, we constructed spliced and unspliced count matrices from our aligned sequencing data using the velocity command-line tool, with GRCh38-p13 as the genome annotation reference. The resulting loom files were processed using scVelo (v0.3.3 - ©Copyright 2024, Theislab)(9). We converted our Seurat object into AnnData format while preserving UMAP coordinates. Preprocessing was performed using scv.pp.filter_and_normalize, selecting genes based on detection and variability. We then applied scVelo's dynamic model (scv.tl.recover_dynamics) to learn the full transcriptional dynamics of splicing kinetics, using a probability-based expectation-maximization framework. This approach estimates cell-specific reaction rates and latent variables, including transcriptional state and internal latent time. RNA velocities were estimated and projected onto UMAP using scv.pl.velocity_embedding_stream, visualizing cellular trajectories as streams. We generated phase portraits of marker genes to illustrate the relationship between spliced and unspliced mRNA abundance. Velocity graphs and pseudotime were calculated to measure the average number of steps required to reach each cell from inferred root cells. To reconstruct lineage relationships and differentiation trajectories, we implemented partition-based graph abstraction (PAGA) using Scanpy's tl.paga function. Edge weights in the PAGA graph represent connectivity confidence, indicating potential differentiation paths. Diffusion pseudotime (DPT) was used to infer a pseudotemporal order of cells along predicted routes, defining a root cell within the starting population. Finally, we extended PAGA with velocity-inferred directionality, providing a graphical map of data topology with weighted edges corresponding to cluster connectivity.

**Proteome Analysis by Mass Spectrometry**

*Sample Preparation*

Protein digestion from $CD44^{+}GLAST^{+}$ and $CD44^{-}GLAST^{-}$ populations of the LGG275 Sample preparation was performed on S-Trap$^{TM}$ micro columns following the manufacturer’s instructions. In brief, protein extracts were diluted in 40 µL of final 5% SDS / 50 mM triethylammonium bicarbonate (TEAB), reduced by 20 mM dithiothreitol (DTT) and held for 10 min at 95°C. Samples were cooled to room temperature and alkylated with 40 mM iodoacetamide (IAA) for 30 min at room temperature in dark. Samples were acidified with phosphoric acid at a final concentration of 1.2% and diluted 6 times in the S-Trap binding buffer (90% methanol / 100 mM TEAB). The resulting protein suspension was transferred to the S-Trap filter via centrifugation at 4000g for 1 min. Trapped proteins were washed three times with a 150 µL S-Trap binding buffer. 1 µg of trypsin in 50 mM TEAB were added to the filter surface and incubated for 2 hours at 47°C. Tryptic peptides were eluted sequentially with 40 µl of 50 mM TEAB / 0.2% aqueous formic acid, and then with 50% of acetonitrinile (ACN) via centrifugation at 4000g. Eluted peptides were vacuum-dried.

*Mass spectrometry*

Experiments were performed on a nano-flow HPLC coupled to a mass spectrometer equipped with a nanoelectrospray source. Peptide samples were solubilized in 0.05% trifluoroacetic acid (TFA) / 2% ACN, and were injected for desalting and pre-concentration on a PepMap®100 C18 precolumn (0.3 mm x 5 mm). Peptide's separation was done on a 25 cm analytical reversed-phase column (75 mm inner diameter)) using a 60 min gradient of 2 to 40% of buffer B (80% ACN, 0.1% formic acid) and a flow rate of 300 nL/min. Spectra were recorded using Xcalibur 4.2 software. MS analyses were performed in data-independent mode (DIA) using 49 windows (13.7 Th) covering a mass range of 361–1033 m/z. Resolution was set to 120,000 for MS1 and 15,000 for MS2. NCE was set to 27%.

*Data analysis*

Spectral data were analyzed via DIANN v1.8.2b27 software [ https://github.com/vdemichev/DiaNN;(10)], using the DIAgui v1.4.2 tool [https://github.com/mgerault/DIAgui;(11)]. For protein databases, we used the Homo sapiens Reference Proteome (UP000005640; release 2025_01; https://www.uniprot.org/) and a homemade contaminant database with the following fixed modification: Carbamidomethylation(C). Data reprocessing, statistical analyses and graphical representations were performed using Perseus v1.6.15.0 software(12).

The mass spectrometry proteomics data have been uploaded to the ProteomeXchange Consortium via the PRIDE partner repository with the dataset identifier PXD064235(13).
